# An ERα-Dependent Hypoxia Response Defines EMT-Adjacent Tumour Regions and Suppresses the Pro-survival Effects of Amiloride in Estrogen Receptor-Positive Breast Cancer

**DOI:** 10.1101/2025.10.03.680339

**Authors:** Jodie R. Malcolm, Jack Stenning, Jacob Pope, Jakub Łukaszonek, Susanna F. Rose, Taylor E. Smith, Lesley Gilbert, Sally R. James, Katherine S. Bridge, William J. Brackenbury, Andrew N. Holding

**Author notes:** Correspondence: Andrew N. Holding. These authors contributed equally (Jodie R. Malcolm, Jack Stenning, Andrew N. Holding).

## Abstract

Estrogen receptor-positive (ER+) breast cancer carries a lifelong risk of recurrence and disease progression, with hypoxia-associated transcriptional signatures linked to poor prognosis and therapy resistance. While the effects of hypoxia on tumour progression are well studied, the impact on ERα epigenomic regulation remains poorly characterised. Here, we demonstrate that activation of hypoxia-inducible factors (HIFs) dramatically remodels ERα chromatin localisation in ER+ breast cancer cells. Transcripts of genes located near hypoxia-induced ERα binding sites are significantly associated with reduced recurrence-free survival in breast cancer patients. Transcriptomic profiling under hypoxic conditions (1% oxygen), with and without ERα depletion by fulvestrant, revealed a hypoxia-induced, ERα-dependent gene expression programme, including upregulation of epithelial sodium channel (ENaC) regulatory subunits that results in acquired sensitivity to the ENaC inhibitor amiloride. Notably, this transcriptional response is spatially correlated with the epithelial-to-mesenchymal hallmark in patient tumours. Our findings establish an interdependence between ERα signalling and the hypoxic response, and present functional evidence that ERα reprogramming offers novel therapeutic opportunities that bypass the need to directly target the hypoxic response.

## Introduction

Estrogen Receptor-α (ERα) drives 70% of breast cancers (Huang *et al*, 2024). Initially, anti-estrogen therapies are effective, but estrogen receptor positive (ER+) disease carries a life-long risk of recurrence (Pan *et al*, 2017; Lindström *et al*, 2018; Yu *et al*, 2019), and ∼30% of patients relapse with metastatic, therapy-resistant disease (Rozeboom *et al*, 2019). Hypoxia, a hallmark of the solid tumour microenvironment, is linked to endocrine resistance and poor outcomes (Li *et al*, 2022; Tutzauer *et al*, 2022; Buffa *et al*, 2010; Place *et al*, 2011; Capatina *et al*, 2024). Although hypoxia-driven epigenetic and transcriptional changes are extensively studied (Collier *et al*, 2023), how hypoxia influences ERα signalling, and conversely, how ERα activity might shape the hypoxic response, remains unresolved. Given the central role of ERα in driving tumour pathophysiology and the established impact of hypoxia on clinical outcomes, we defined the relationship between the two signalling pathways.

The hypoxic transcriptional response is primarily governed by the Hypoxia-Inducible Factors (HIFs). Of the three α-subunits (HIF-1α, HIF-2α, HIF-3α), HIF-1α and HIF-2α are best characterised; HIF-3α is less defined (Gu *et al*, 1998; Duan, 2016; Ortmann, 2024). Under normoxic conditions, HIF-1α and HIF-2α are hydroxylated by the oxygen-dependent prolyl hydroxylases PHD1–3 (Tanimoto *et al*, 2000; Schofield & Ratcliffe, 2004). These enzymes target proline pairs on HIF-1α and HIF-2α, recruiting the von Hippel-Lindau (pVHL) E3 ubiquitin ligase, triggering degradation.

At low oxygen, PHD activity is inhibited, allowing HIF-1α and HIF-2α to stabilise and heterodimerise with the constitutive HIF-1β/ARNT. These heterodimers, named HIF-1 and HIF-2, translocate to the nucleus and activate transcriptional signalling. Targets include *VEGFA, GLUT1* and *GAPDH,* which promote angiogenesis and modulate glucose uptake and metabolism (Forsythe *et al*, 1996; Muz *et al*, 2015; Chen *et al*, 2001; Hayashi *et al*, 2004; Higashimura *et al*, 2011).

A second layer of regulation is provided by Factor Inhibiting HIF (FIH), which hydroxylates an asparagine on HIF-1α and HIF-2α, blocking interaction with the co-activator p300 (Lando *et al*, 2002) and reducing HIF transcriptional activity. As FIH is more active in moderate hypoxia than the PHDs, partially active HIF proteins accumulate before full activation (Figg *et al*, 2021; Wang *et al*, 2025).

Despite shared regulatory mechanisms, HIF-1α and HIF-2α act on distinct timescales. HIF-1α stabilises in acute hypoxia (4-16 hours), whereas HIF-2α dominates in chronic hypoxia (>24 hours). Both are relevant in breast cancer, where hyperproliferation and disorganised vasculature leads to a dynamic and irregular oxygen landscape across the tumour microenvironment (Liu *et al*, 2022a; Koh & Powis, 2012).

Beyond oxygen sensing, HIFs are also subject to hypoxia-independent regulation. Examples of this in normal and malignant physiology include cytokine-driven upregulation in the placenta (Iriyama *et al*, 2015), insulin-mediated activation (Zelzer *et al*, 1998), MAPK inhibition of nuclear export (Mylonis *et al*, 2006) and oncogene-driven kinase-mediated activation in in myeloproliferative neoplasms and prostate cancers (Kealy *et al*, 2024; Casillas *et al*, 2021).

HIF-1α–ERα crosstalk links hypoxic and estrogen responses (Wolff *et al*, 2017; George *et al*, 2012), and promotes endocrine resistance (Morotti *et al*, 2019; Yang *et al*, 2010, 2015; Capatina *et al*, 2024). Similar roles have been proposed for HIF-2α (Fuady *et al*, 2016). In parallel, HIF-1β, as ARNT, directly interacts with the ERα (Brunnberg *et al*, 2003; Matthews & Gustafsson, 2006). Moreover, ligand activation of the aryl hydrocarbon receptor (AhR), which heterodimerises with ARNT, modulates ERα binding (Matthews *et al*, 2005) and transcription (Madak-Erdogan & Katzenellenbogen, 2012).

Oxygen availability is also critical for chromatin regulation, with histone demethylases acting as key oxygen sensors (Melvin & Rocha, 2012; Collier *et al*, 2023). Lysine-specific demethylase 1 (LSD1) (Shi *et al*, 2004), removes H3K4 mono- and dimethylated marks via an oxygen- and FAD-dependant mechanism (Culhane & Cole, 2007; Perillo *et al*, 2020) coupling chromatin to oxygen availability. LSD1 also recruits CoREST to influence broader epigenetic regulation (Song *et al*, 2020). A second family, the Jumonji C (JmjC) domain demethylases (Tsukada *et al*, 2006), act on multiple H3 lysines (Seward *et al*, 2007; Shin & Janknecht, 2007) and require molecular oxygen. As they share structural features FIH (Loenarz & Schofield, 2011) and PHD1-3, JmjC-domain demethylases are also inhibited by dimethyloxalylglycine (DMOG), a common hypoxia mimetic (Chan *et al*, 2016; Hopkinson *et al*, 2013).

The chromatin landscape plays a key role in regulating ERα transcriptional activity (Joseph *et al*, 2010). In MCF7 cells, KDM6B shows oxygen sensitivity (Prickaerts *et al*, 2016) and participates in an ERα/KDM6B regulatory axis (Liu *et al*, 2022b), demonstrating oxygen-dependent demethylases modulate ERα. Reciprocally, ERα has been shown to recruit LSD1 to target genes (*TFF1*, *GREB1, CTSD*, and *MYC*), and their expression is abolished upon LSD1 knockdown (Garcia-Bassets *et al*, 2007). Our own qPLEX-RIME data detected LSD1 (KDM1A) and KDM6A in ERα complexes following estrogen stimulation (Zhao *et al*, 2024), and LSD1 recruitment generates localised reactive oxygen species associated with ERα transcriptional activity (Perillo *et al*, 2014).

Here, we investigate the interdependence of hypoxic and estrogen signalling. Genome-wide profiling in MCF7 cells revealed ERα cistrome reprogramming in response to DMOG, with gained binding sites enriched for ERE, FOXA1, and AP-1 motifs. Transcriptomic analysis of hypoxic (1% oxygen) MCF7 and T47D cells, with and without ERα degradation by fulvestrant, defined a distinct ERα-dependent hypoxic response (EDHR). EDHR genes were the regulatory subunits of the epithelial sodium channel (ENaC), whose altered expression depended on both hypoxia and active estrogen signalling. Spatial transcriptomics further revealed significant co-localisation of the EDHR signature epithelial-to-mesenchymal transition regions in ER+ tumours. Moreover, the ENaC inhibitor amiloride, which increases MCF7 cell viability in normoxia (Ware *et al*, 2021), had the opposite effect under hypoxic conditions, reducing cell viability.

Together, these results define a novel ERα-dependent hypoxia response, and identify ENaC as a potential therapeutic vulnerability in hypoxic ER+ disease. To our knowledge, this is the first characterisation of a ERα-dependent hypoxia response, revealing a previously unrecognised layer of hormonal control over the hypoxic breast cancer transcriptome.

## Materials and Methods

### Cell Culture

MCF7 and T47D cells were routinely cultured in Dulbecco’s Modified Eagle Medium (DMEM) with high glucose, L-glutamine, phenol red and sodium pyruvate (Gibco, #11995-073), supplemented with 10% fetal bovine serum (FBS; Gibco, #A5256801), unless stated otherwise. Cell lines were maintained in a humidified CO_2_ incubator (LEEC, Precision 190D) at 37 °C, 5% CO_2_ and atmospheric O_2_ (normoxia) in 2D. All cells were authenticated by STR genotyping (Masters *et al*, 2001). Cell lines were confirmed *Mycoplasma-*free by immunofluorescent visualisation of contaminating *Mycoplasma* DNA with DAPI (Uphoff *et al*, 1992) and by PCR analysis for mycoplasma DNA (Siegl *et al*, 2023). For hypoxic experiments, breast cancer cells were incubated in a humidified Baker Ruskinn InvivO2 oxygen workstation at 37 °C, 5% CO_2_ and 1% O_2_ for up to 48 hours.

### Cell Lysis and Western Blot

Cells were seeded in a 6-well tissue culture-treated plate with DMEM supplemented with 5% FBS (Gibco), and incubated at 37 °C and 5% CO_2_ for 24 hours. Next, cells were either maintained in atmospheric O_2_ or moved into the hypoxic InvivO_2_ workstation for 8 or 48 hours. At experimental endpoint, cells were washed with ice-cold phosphate-buffered saline (PBS) and lysed in RIPA buffer (Merck) with cOmplete^TM^ EDTA-free protease inhibitor cocktail (Sigma-Aldrich) and PhosSTOP^TM^ phosphatase inhibitor (Roche). Samples were left to sit on ice for 10 minutes and then centrifuged to pellet cellular debris (16,000 g, 10 minutes, 4 °C). Supernatant was collected for western blot analysis and the pellet was discarded. Cell lysate was prepared 1:4 (v/v) in Laemmli Sample Buffer (Bio-Rad) with β-mercaptoethanol (1:9 v/v) and boiled at 100 °C for 15 minutes. Samples were quenched on ice, briefly centrifuged and loaded onto a 10% SDS-polyacrylamide gel. Samples were loaded into the stacking gel by applying 70 V for 15 minutes and resolved at 120 V until sample dye was at the bottom of the gel. Next, gels were loaded onto Trans-Blot Turbo Mini polyvinylidene difluoride (PVDF) membranes (Bio-Rad) and transferred in a TransBlot Turbo Transfer Device (Bio-Rad) using a default 1.5 mm gel programme of 25 V for 10 minutes. PVDF membranes were blocked at room temperature for 1 hour in a blocking solution consisting of 4% (w/v) non-fat dairy milk (Marvel) in Tris-buffered saline (TBS) with 0.1% (v/v) Tween 20 (TBST). Membranes were washed three times in TBST (5 minutes per wash) before incubating overnight at 4°C in anti-HIF-1α (Cell Signalling, 36169; 1:1,000) or anti-α-Tubulin (Merck, T6199; 1:5,000) antibodies prepared in blocking solution. Membranes were washed three times in TBST (5 minutes per wash) and incubated at room temperature for one hour in anti-rabbit (HIF-1α; Cell Signalling, 7074S; 1:10,000) or anti-mouse (α-Tubulin; Cell Signalling, 7076; 1:10,000) IgG HRP-linked antibody in blocking solution. Membranes were then washed three times in TBST (5 minutes per wash). Next, membranes were incubated for 5 minutes in western blotting luminol reagent (Santa Cruz) and imaged on an iBright FL500 imager (Thermo Fisher Scientific).

### RNA-Seq

Breast cancer cells were seeded into 10 cm dishes at 6,500 cells/cm^2^ and incubated at 37°C and 5% CO_2_ for 24 hours. Next, cells were treated with either 100 nM of fulvestrant (Cayman) or ethanol vehicle control and returned to a humidified CO_2_ incubator in atmospheric O_2_. After 48 hours, dishes were either left in these normoxic conditions or transferred to the hypoxic InvivO_2_ workstation for a further 48 hours. At the end of the incubation period, cells were rinsed in PBS and trypsinised for 5 minutes at 37°C, after which the trypsin was neutralised with culture medium and cells were centrifuged at 300 g for 5 minutes. The supernatant was removed and the cells were washed in PBS and centrifuged again at 500 × *g* for 5 minutes. The supernatant was removed and the pellet of cells were resuspended in PBS. For ultra-low temperature storage of samples before RNA isolation, RNAlater (Sigma) was added before storing at −80°C. RNA was prepared from the cells using the Monarch Total RNA mini kit (New England Biolabs) following the manufacturer’s instructions. The purified RNA samples were submitted to the University of York Bioscience Technology Facility (BTF) Genomics Lab for library preparation. Libraries were sequenced on an Illumina NovaSeq 6000 platform (Azenta Life Sciences) using a 2 × 150 bp paired-end configuration with a minimum sequencing coverage requirement of 20 million reads per sample. Base calling was performed using Illumina’s real-time analysis software. For each cell line and each experimental condition, a minimum of three biological replicates (n = 3) were prepared for sequencing.

### Chromatin Immunoprecipitation Followed by High-Throughput Sequencing (ChIP-Seq)

MCF7 cells were seeded onto 15 cm tissue culture-grade dishes and cultured as described above. After 24 hours, breast cancer cells were treated with 2 mM DMOG or vehicle control (dH_2_O) for 16 hours (Schödel *et al*, 2011). DMOG is a broad-spectrum 2-oxoglutarate dioxygenase inhibitor that has previously been demonstrated to provide a MCF7 transcriptional response more similar to hypoxia than the use of selective PHD inhibitors (Chen *et al*, 2022). ChIP-seq was performed as previously described (Holding *et al*, 2018). A total of 10 μg of ERα antibody (Abcam, ab3575) was used per ERα ChIP (Glont *et al*, 2019). Samples were submitted to the University of York Bioscience Technology Facility (BTF) Genomics Lab for library preparation and Illumina short read sequencing. Libraries were sequenced on an Illumina NovaSeq 6000 platform (Novogene) with 2 × 150 bp paired-end configuration with a minimum sequencing coverage requirement of 20 million reads per sample. Sequencing reads were aligned to the GRCh38/hg38 human reference genome using BWA (v1.11). Reads that mapped to genomic loci listed in the ENCODE blacklist for hg38 (hg38-blacklist.v2.bed) were removed to minimise artifacts (Amemiya *et al*, 2019). Peaks were called using MACS2 (2.1.4) (Zhang *et al*, 2008) and compared using BEDtools (v2.28.0). Motif analysis was undertaken with Homer (v4.11) against the inbuilt list of known motifs. For visualisation and comparative analysis, bamCoverage, ComputeMatrix and plotHeatmap packages were used from deepTools (v3.5.6) (Ramírez *et al*, 2016).

### Viability Assay

Cells were seeded at 40,000 cells per well in 24-well tissue culture-treated plates in 500 µL DMEM supplemented with 5% FBS and allowed to adhere in normoxic conditions. After 24 hours, cells were treated with amiloride hydrochloride (Merck, A7410). Working concentrations were freshly prepared by serial dilution from a 100 mM stock solution in DMSO to final concentrations of 1 nM, 10 nM, 100 nM, 1 µM, 10 µM, and 100 µM, maintaining a constant DMSO vehicle concentration across all treatment groups, including the vehicle control. Plates were then returned to either normoxic (21% O₂) or hypoxic (1% O₂) culture conditions for 72 hours.

At the experimental endpoint, media was aspirated, and cells were washed once with phosphate-buffered saline (PBS). Cell viability was assessed by staining with 7-Aminoactinomycin D (7-AAD; BioLegend, 420404) at a 1:1000 dilution in FACS buffer (PBS supplemented with 2% FBS and 5 µM EDTA). Cells were incubated with 200 µL of the 7-AAD staining solution per well for 10 minutes on ice in the dark, followed by a wash with PBS. Adherent cells were detached using 2x trypsin (100 µL per well; Gibco) for 2-3 minutes at 37 °C, then neutralized with 100 µL PBS + 2% FBS. Cell suspensions were transferred into 96-well V-bottom plates for flow cytometric analysis.

Flow cytometry was performed on a CytoFLEX LX (Beckman Coulter). Cells were gated based on forward and side scatter properties to exclude debris and doublets, and viability was determined by exclusion of 7-AAD positive cells. Data were analyzed using FlowJo software. Viability was expressed as the percentage of 7-AAD negative (live) cells.

### Spatial Transcriptomics

Preprocessed spatial transcriptomics data were obtained from the supplementary materials accompanying the *SpottedPy* study (Withnell & Secrier, 2024). All code used for reanalysis is available through the Data availability section.

### Public Database Searches

To investigate chromatin accessibility and gene expression in breast cancer, publicly available datasets were analysed in addition to those generated in this study. ATAC-seq data were obtained from the Gene Expression Omnibus under accession number GSE133327 (Miar *et al*, 2020). Transcriptomic data (Godet *et al*, 2019) were retrieved from GEO for differential expression analysis under accession number GSE111653.

For metagene survival analysis, the *survminer* R package (v0.4.9) was used to determine the optimal expression cut point using the surv_cutpoint function and was applied to expression data from The Cancer Genome Atlas (TCGA) Breast Cancer Patient Cohort (Cancer Genome Atlas Network, 2012)

For individual gene survival analysis, Kaplan–Meier survival analyses were performed using KMplot (Lánczky & Győrffy, 2021; Győrffy, 2024). For *SCNN1B*, recurrence-free survival (RFS) was assessed in ERα-positive patients (defined by immunohistochemistry), using auto-selected percentile cut-off and including all follow-up times (n = 2,633). The same parameters were used for *SCNN1G* (n = 947); the lower sample number reflects the limited availability of *SCNN1G* expression data across all included datasets. For *SCNN1A*, survival analysis was conducted using the same parameters for both RFS (n = 4,929; ERα+ = 2,633; ERα- = 2,651) and overall survival (n = 1,879; ERα+ = 754; ERα- = 520); status was stratified on the basis of immunohistochemistry.

## Results

### Inhibition of Dioxygenases by the Hypoxia Mimetic DMOG Redistributes ERα Binding across the Genome of ERα+ Breast Cancer Cells

To investigate the impact of HIF stabilisation on ERα chromatin occupancy, we performed ChIP-seq in MCF7 breast cancer cells treated with either the hypoxia mimetic DMOG or vehicle control, following conditions previously established for HIF ChIP-seq (Schödel *et al*, 2011).

Our analysis identified substantial changes in the genomic locations occupied by ERα in response to DMOG, which we classified into three categories of ERα binding: (i) conserved sites, where ERα binding was detected in both DMOG- and vehicle-treated cells over input control (**Supplementary Table S1**); (ii) lost sites, where ERα binding was no longer identified upon dioxygenase inhibition (**Supplementary Table S2**); and (iii) gained sites, where ERα occupancy was only identified after the DMOG treatment (**Fig. 1a**, **Supplementary Table S3**).

**Figure 1.**
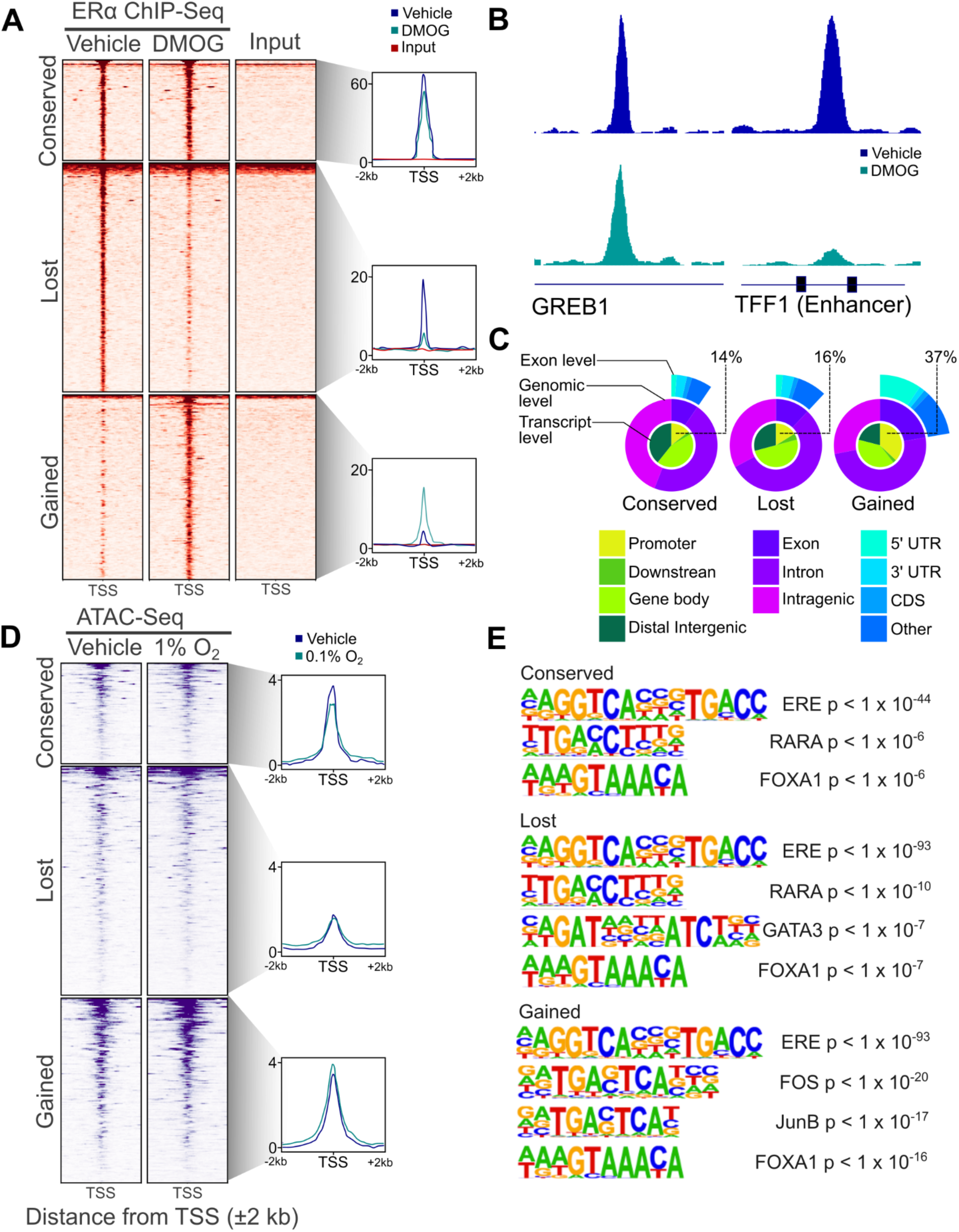
DMOG treatment leads to an epigenomic reprogramming of ERα. (A) ERα ChIP-seq Heatmap for ERα binding sites demonstrates a redistribution of ERα at the majority of binding sites in response to DMOG. (B) Individual analysis of *GREB1* promoter [GRCh38 chr2:11,539,400-11,539,900] and *TFF1* enhancer loci [GRCh38 chr21:42,376,000-42,377,000] show that classic estrogen response genes have differing responses to DMOG. (C) Analysis of conserved, gained, and lost ERα ChIP-seq loci shows that the gained sites are found biased towards promoter regions (37%) when compared to the other conserved and lost ERα binding sites. (CDS = Coding Sequence). (D) ATAC-Seq (GSE133327) analysis of identified ERα binding sites shows no identifiable change in chromatin accessibility in response 48h at 0.1% oxygen. (E) Top identified motifs for the conserved and lost sites included the ERE and common ERα binding partners and typical of the ERα ChIP-seq. Analysis of gained sites identified ERE and an increased presence of the FOS (22.66%, rank #2, p < 1 x 10^-20^) and JunB (25.78%, rank #3, p < 1 x 10^-17^) motifs. In contrast, the JunB motif was not identified at either the lost or conserved sites, and while FOS was detected at lost sites, it was found at a much lower frequency (7.01%, rank #45, p = 0.01).

At the *GREB1* gene locus, a well-studied ERα target gene and transcriptional cofactor, ERα binding was conserved at both the promoter and enhancer across conditions. In contrast, ERα binding was markedly reduced in response to DMOG treatment at both the promoter and enhancer for TFF1, another classical estrogen-responsive gene (**Fig. 1b**). Together, these findings illustrate divergent responses at established ERα-bound regulatory regions of estrogen-responsive genes (Rae *et al*, 2005; Ribieras *et al*, 1998). Additionally, we provide an example of a gained ERα binding event at the promoter of *CASC4*, also known as *GOLM2,* which to the best of our knowledge has not previously been reported as an estrogen regulated gene (**Supplementary Fig. S1a**).

To explore how these changes in ERα binding might contribute to genomic function, and to provide confidence that these results were not artefactual, we performed genomic feature annotation of our ERα ChIP-seq peaks. This analysis revealed that gained binding sites were disproportionately enriched at promoter regions (37%) compared to conserved (14%) and lost (16%) sites (**Fig. 1c, Supplementary Fig. S1b**), consistent with a role of direct ERα-mediated transcriptional regulation.

As hypoxia and HIFs are closely associated with changes in chromatin accessibility across a range of cell types (Batie *et al*, 2022; Xin *et al*, 2020; Wang *et al*, 2020), we hypothesised that the redistribution of ERα might be driven by significant alterations in chromatin accessibility at gained and lost ERα sites. To test our hypothesis, we analysed published ATAC-seq generated from MCF7 cells exposed to 48 hours of 0.1% oxygen (GSE133327) (Lombardi *et al*, 2022) at the genomic loci corresponding to our ERα binding categories (**Fig. 1a**). The resulting ATAC-seq read density plots showed no changes in chromatin accessibility that could account for the scale of ERα binding alterations observed, suggesting that the redistribution of ERα reflects mechanisms distinct from changes in chromatin accessibility (**Fig. 1d**).

Motif analysis of conserved, lost and gained ERα binding loci revealed Estrogen Response Elements (EREs) as the most significantly enriched motif across all three classes. Motifs found within the top 10 enriched for each class, after filtering for redundant results, are shown in **Fig. 1e** (unfiltered data is provided in **Supplementary Fig. S2**). Beyond the ERE, all classes showed significant enrichment of the binding motif for FOXA1, an established ERα pioneer factor essential for ERα activity (Hurtado *et al*, 2011) and a known HIF-2α interactor (Fu *et al*, 2019). Gained sites were also enriched for FOS and JunB motifs, whereas conserved and lost sites featured RARα, a known ER-dependent cooperative factor (Ross-Innes *et al*, 2010). GATA3 was only identified as among the top motifs for lost sites. The elevated ranking of Fos and Jun motifs at gained ERα sites is concomitant with the role of AP-1 interacting with ERα at these loci and with AP-1 being a known HIF-1 interactor (Laderoute, 2005). The Hypoxia Response Element (HRE) was not enriched in the motif analysis in any of the ERα binding classes.

Taken together, these results demonstrate that the inhibition of α-ketoglutarate-dependent dioxygenases, mimicking hypoxia, leads to extensive redistribution of ERα binding across the genome in ER+ breast cancer cells.

### Transcripts of Genes Associated with the Redistribution of ERα in Response to DMOG Predict Worse Patient Outcome

Having identified distinct classes of ERα binding sites altered by DMOG treatment, we next investigated the relationship between transcripts associated with these epigenomic changes and breast cancer subtype. We hypothesized that the expression of transcripts associated with gene proximal to these gained and lost ERα binding sites would be predominantly associated with ER+ breast cancer cell line models and correlate with clinical outcomes. To strengthen these findings, we leveraged publicly available RNA-seq data from breast cancer cell lines exposed to 1% oxygen, which allowed us to confirm that these transcriptional changes represent authentic hypoxic responses validating that the effects observed with the hypoxic mimetic DMOG are consistent with *bona fide* hypoxia.

Given that ERα binding at promoters and enhancers correlates with estrogen-induced expression of neighbouring genes (Carroll *et al*, 2005), we mapped the ‘lost’ and ‘gained’ classes of ERα binding events to proximal genes. Our mapping enabled the construction of two ERα-associated gene networks, corresponding to our ‘lost’ and ‘gained’ sites. From these, we derived the two corresponding transcriptional signatures to integrate our ChIP-seq data with RNA-seq profiles and to assess their relevance in published breast cancer datasets. This approach has previously been used successfully to link ERα chromatin occupancy to downstream gene expression and clinical outcome in breast cancer (Ross-Innes *et al*, 2012).

Principal Component Analysis (PCA) of these gene signatures was applied to a publicly available transcriptional dataset of 31 breast cancer cell lines exposed to either normoxia or hypoxia (GSE111653, (Godet *et al*, 2019)). Clustering on the subset of transcripts within our ERα binding loss-associated signature demonstrated strong subtype-specific separation, with all ERα+ breast cancer cell lines, including ER+/HER2+ cells, clustering separately (PC1 > 0) from cell lines representing all other breast epithelial and breast cancer subtypes (**Fig. 2a**). In contrast, TNBC and other ERα-cell lines demonstrated much more limited variance captured by PC1 and PC2.

**Figure 2.**
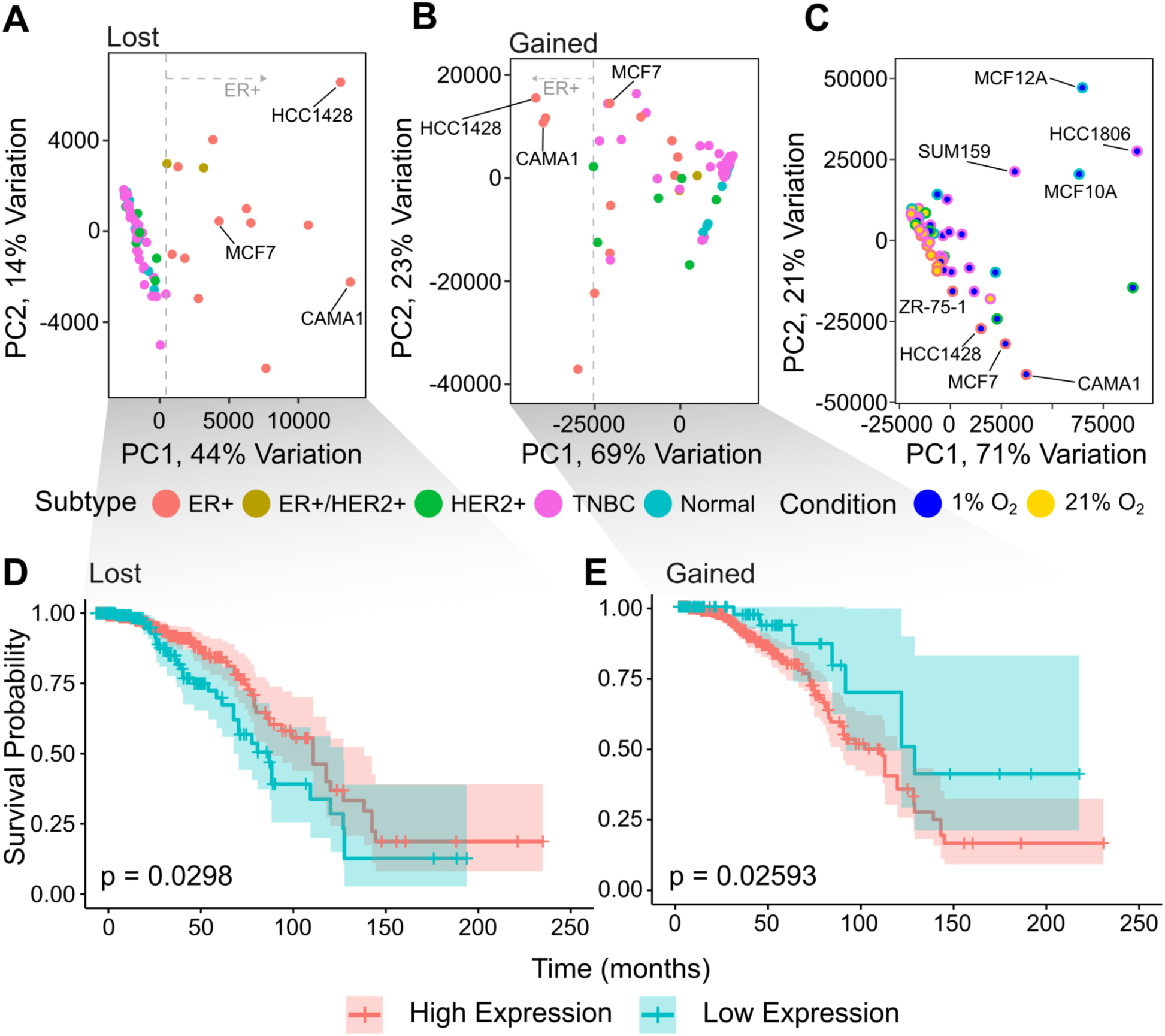
Genes proximal to changes in ERα binding are associated with breast cancer cell lines and patient survival. (A) Principal Component Analysis (PCA) of transcripts associated with ERα binding loss in RNA-seq data from 32 breast cancer cell lines (GSE111653) identified that 41% of the variance was explained by PC1; and that for values greater than 0, PC1 aligned with ERα+ subtypes. (B) PCA of transcripts associated with gained ERα sites showed an alignment of negative PC1 values with an increasing proportion of ERα+ cell lines. (C) PCA performed on ER+ hypoxia-responsive transcripts as established by differential analysis of the ER+ cell lines. (D) Survival analysis of the TCGA cohort of the loss-associated transcriptional signature shows that decreased expression of these genes is significantly associated with worse RFS. (E) Survival analysis of the TCGA cohort of the gain-associated transcriptional signature shows that increased expression of these genes is significantly associated with worse RFS.

Clustering on the subset of transcripts in our ERα binding gain-associated signature associated ERα+ and HER2+ cell lines with lower values of PC1. While not displaying as distinct separation as our loss-associated signature, all ERα+ cell lines have a PC1 value < 0, while the majority of TNBC samples clustered together with a PC1 value > 0 (**Fig. 2b**).

For comparison, PCA was performed using hypoxia-responsive transcripts as established by differential expression analysis of the ERα+ cell lines from the same dataset. The PCA captured variance associated with oxygen levels, demonstrating separation by condition. However, unlike our ChIP-seq derived signatures, PC1 no longer captured variance associated with ERα+ cell lines, instead describing features associated with the normal cell lines (MCF12A, MCF10A) and the TNBC line (HCC1806). In contrast, the hypoxic response in ERα+ cell lines such as CAMA1, HCC1428, and MCF7, was now mostly represented along PC2 **(Fig. 2c)** and contributing less variance. These results indicate that our ChIP-seq-derived signatures more specifically captures ERα-associated transcripts in comparison to the hypoxia differential expression profile derived from the ERα+ cell lines within the data set.

To contextualise the potential significance of our hypoxia mimetic-induced ERα binding events in breast cancer outcome, we stratified The Cancer Genome Atlas (TCGA) Breast Cancer Patient Cohort (Cancer Genome Atlas Network, 2012) on the basis of the expression of each of the ERα gene transcription signatures. Kaplan-Meier analysis of recurrence-free survival (RFS) revealed a significantly better outcome for patients with a high expression of transcripts within our loss-associated signature (p = 0.0298, log-rank test, n = 691, **Fig. 2d**), whereas RFS was significantly reduced for patients with high expression of the transcriptional signature from gain-associated signature (p = 0.0259, log-rank test, n = 691, **Fig. 2e**). In ERα-tumours, neither signature showed a significant association with outcome (data not shown).

To further investigate if the hypoxia response observed was specific to ERα+ breast cancer, we next compared our ERα+ gene signatures against an established generalised hypoxia metagene, which has previously been shown to be highly prognostic for multiple cancers (Buffa *et al*, 2010). No overlap was observed between our ChIP-seq-derived gene signatures and the Buffa hypoxia metagene, supporting our hypothesis that the gene signature we see is ERα+ breast cancer specific. Similarly, we detected no overlap with the 48-gene pan-cancer signature described by Lombardi *et al*. (2022), further supporting that the signature we identified is specific to ERα+ breast cancer.

These results demonstrate that: (i) hypoxia induces specific gene expression changes in ERα+ cell lines associated with our ChIP-seq-derived networks; (ii) our ChIP-seq-derived networks are predictive of patient recurrence-free survival; and (iii) the genomic redistribution of ERα in response to DMOG, and the resulting HIF stabilisation, define a transcriptional programme that is distinct from the hypoxic response in TNBC cell lines.

### Transcriptomic Analysis of ERα+ Breast Cancer Cells show Expected Response to Hypoxia and a Targeted Antiestrogen Regiment

To investigate the interaction between ERα- and hypoxia-mediated signalling in ERα-positive breast cancer, we designed a multifactorial experiment to enable transcriptomic profiling of MCF7 and T47D cell lines following perturbations in ERα protein expression and oxygen availability.

Our ChIP-seq experiments employed the hypoxia mimetic DMOG, applying timescales established in previous HIF ChIP-seq studies (Schödel *et al*, 2011; Lombardi *et al*, 2022). However, RNA-seq requires substantially fewer cells, enabling us to culture the cells in tumour-relevant hypoxia (1% O₂) for the time course of the experiment. Since HIF stabilisation in response to hypoxia has been shown to vary significantly between cell types and culture conditions (Bracken *et al*, 2006), we initially characterised HIF-1α protein dynamics in MCF7 and T47D cells. Immunoblotting revealed a robust induction of HIF-1α protein after 8 hours in hypoxia, which was substantially reduced at 48 hours (**Fig. 3a left, Supplementary Fig. S3**). Guided by our immunoblot data, and prior evidence demonstrating that the transcripts of classical HIF target genes (*EGFA*, *CA9*, *PDK1*, *BNIP3*, and *BNIP3L)* (Rogers *et al*, 2024) and the ERα/HIF-1α co-regulated gene *SLC38A2 (Morotti et al, 2019)* peaked at 48 hours in 1% oxygen, the experiment duration was extended accordingly. We then confirmed fulvestrant-mediated degradation of ERα over this timeframe by western blot, showing near-complete loss by 48 hours, and undetectable levels of ERα protein at 96 hours in MCF7 cells under hypoxic conditions (**Supplementary Fig. S4**).

**Figure 3.**
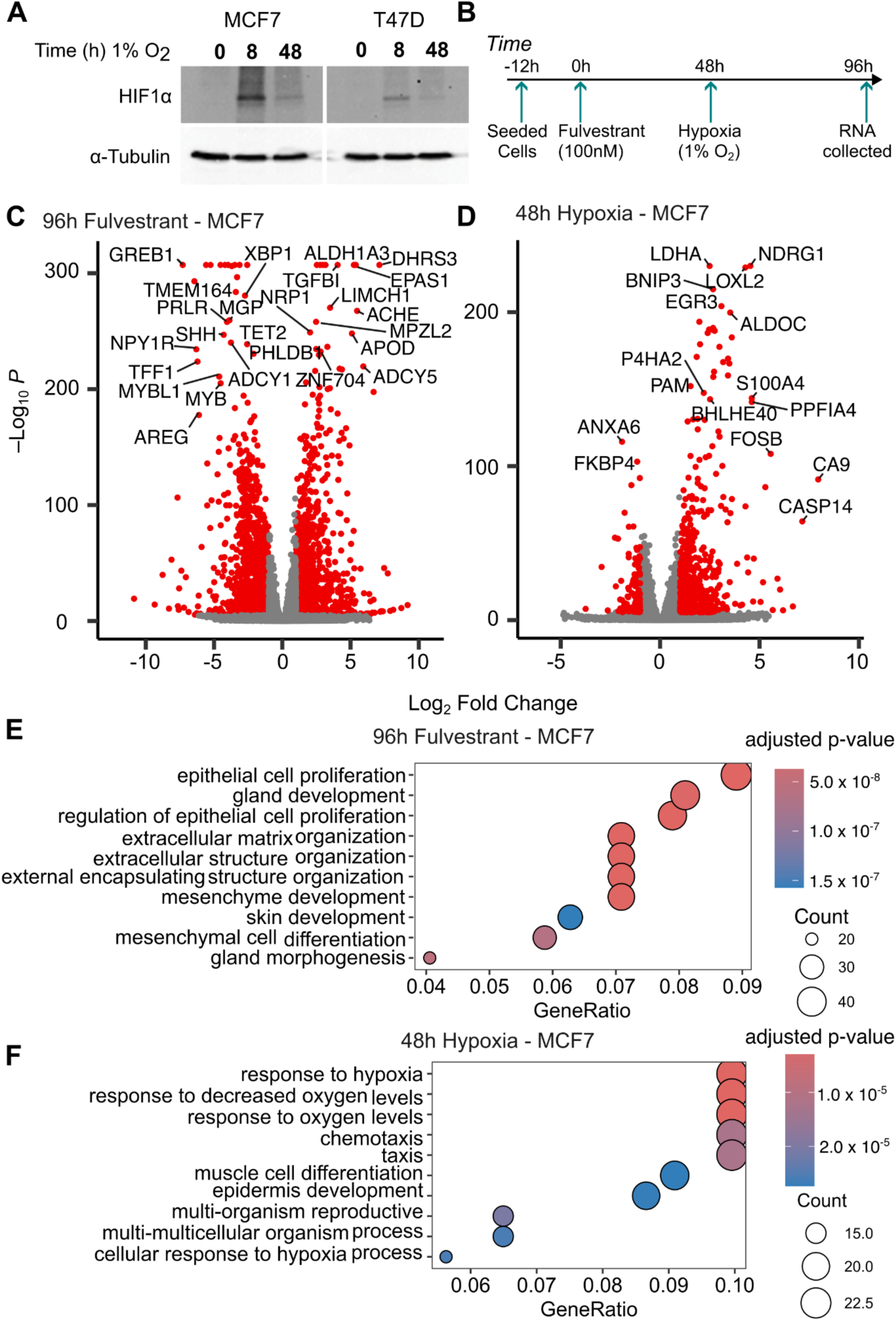
RNA-seq of 96h response to 100 nM fulvestrant and 48 hour response to hypoxia aligns with previously reported responses. (A) Western blots of MCF7 and T47D cell lines for HIF-1α demonstrated an increase in HIF-1α expression at 8 hours of hypoxic culture, which then decreased at 48 hours. (B) Scheme of experimental time course. (C) 96 hour treatment with fulvestrant led to a loss in expression of known ERα target genes, e.g. *GREB1* and *TFF1*. (D) 48 hours of hypoxia (1% oxygen) lead to induction in the expression of known HIF target genes, e.g. *CA9*. (E) GO enrichment analysis of differentially expressed genes (DEGs) in response to fulvestrant treatment demonstrated a significant association with estrogen-related biological processes. (F) GO enrichment analysis of DEGs associated with 1% oxygen demonstrated a significant association of our data with known biological processes.

Based on these findings, cells were treated with 100 nM fulvestrant or vehicle for 48 hours, followed by a further 48-hour incubation at 1% oxygen prior to RNA collection (**Fig. 3b**). Our experimental design enabled interrogation of ERα and hypoxia responses both independently and in combination, allowing comparison of: (i) the fulvestrant response in normoxia, to define ERα-dependent transcription; (ii) the hypoxic response in the absence of fulvestrant, to define hypoxia-dependent transcription; and (iii) the hypoxia response in the absence of ERα, to define ERα-dependant hypoxia transcription.

Differential gene expression analysis of MCF7 cells treated with fulvestrant under normoxia showed significant reduction in mRNA expression of well-characterised ERα-regulated genes, including *GREB1* and *TFF1* (*Padj* < 0.001; **Fig. 3c, Supplementary Table S4**) (Mohammed *et al*, 2013; Holding *et al*, 2018; Baron *et al*, 2007; Spadazzi *et al*, 2021; Hodgkinson *et al*, 2018). Notably, ERα degradation corresponded to a significant increase in *EPAS1* (HIF-2α) levels in MCF7 (**Fig. 3c**) and T47D cells (**Supplementary Fig. S5, Supplementary Table S5**), in line with previous findings (Fuady *et al*, 2016). Analysis of gene expression alterations in hypoxic cells revealed significant upregulation of *bona fide* hypoxia-responders in MCF7 (**Fig. 3d, Supplementary Table S6**) and T47D (**Supplementary Fig. S5, Supplementary Table S7**) cells, including *CA9, FOSB* and *VEGFA*, consistent with known hypoxia-induced gene expression profiles *(Forsythe et al, 1996; Wykoff et al, 2000; Turner et al, 2002; Premkumar et al, 2000)*. Gene Ontology (GO) analysis of biological processes (BP) enriched following fulvestrant treatment of MCF7 (**Fig. 3e**) and T47D cells (**Supplementary Fig. S5**) demonstrated a significant negative enrichment for processes involved in regulation of cell growth, cell proliferation and extracellular matrix (ECM) structure and organisation (adjusted p < 0.05). BPs enriched in hypoxic MCF7 (**Fig. 3f**) and T47D cells (**Supplementary Fig. S5**) were positively enriched for responses to hypoxia, decreased oxygen levels and changes in glucose metabolism (adjusted p < 0.05).

Together, these results align with previously characterised individual transcriptional programmes of ERα and hypoxia, and validate the robustness of our RNA-seq dataset for further investigation into ERα–hypoxia interplay in breast cancer.

### ERα Loss Blunts and Alters Coordination of the Hypoxic Transcriptional Response

To investigate the contributions of ERα and hypoxia to each other’s transcriptional programmes, we stratified differentially expressed genes (DEGs) based on their significant response to hypoxia, fulvestrant-mediated ERα degradation, or both. Across all conditions, we identified 11,622 DEGs (adjusted p < 0.05). Of these, 7,093 genes significantly altered in hypoxia, with 2,344 transcripts (20%) uniquely responsive to hypoxia alone (our ‘Hypoxia’ gene set). Fulvestrant treatment resulted in 9,278 DEGs, of which 4,529 (39%) were specific to ERα loss (our ‘ERα targets’ gene set). Notably, 4,749 genes (41%) were differentially regulated under both conditions, representing an overlap of both responses (our ‘Shared Response’ gene set). To determine whether the overlap between ERα-regulated and hypoxia-regulated genes was greater than expected by chance, we performed a hypergeometric test on the overlap of both responses using all expressed genes as the background. The observed intersection was statistically significant (p < 0.001), supporting a non-random, functionally relevant convergence of responses between the two pathways (**Fig. 4a**).

**Figure 4.**
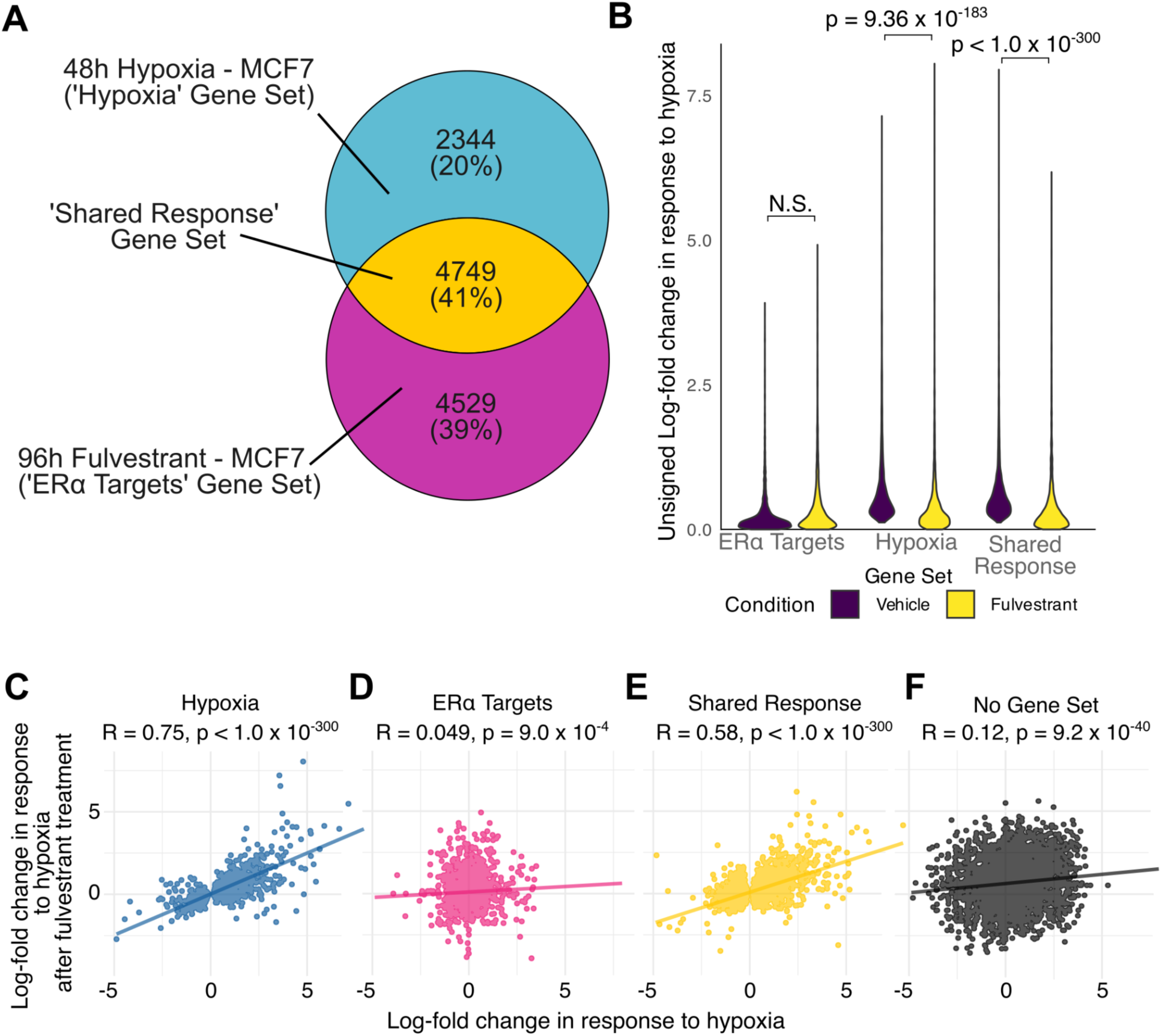
Target ERα signalling significantly dampens the HIF response on MCF7 cells. (A) Using the transcriptomic data from Figure 3, we stratified transcriptional responses into those that are unique to a ‘Hypoxia’ gene set or an ‘ERα Target’ gene set, or those that were shared between the two conditions as a ‘Shared Response’ gene set. (B) Analysis of the gene sets demonstrated that ERα-specific targets were not responsive to hypoxia. Our hypoxic gene set showed significant reduction in the absolute LFC hypoxic response of these transcripts in response to fulvestrant. Our shared response showed both a reduction in the maximum absolute LFC and a greater effect size in the reduction of the transcriptional response, indicated by the reduced p-value, in response to fulvestrant. (C) LFC for our hypoxia genes set showed strong correlation in the hypoxia response with ERα and the hypoxic response after fulvestrant is used to degrade ERα. (D) Our ERα gene set showed very weak correlation in the hypoxic response between conditions (E) Our shared response showed a moderate correlation in hypoxic response between conditions. (F) Genes not within our genes showed very weak correlation between conditions.

We next examined the absolute Log_2_ Fold Change (LFC) in response to hypoxia in MCF7 cells, with or without the ERα (i.e. following vehicle or fulvestrant treatment) for each gene set. To avoid confounding effects, genes within the ‘Shared Response’ gene set were excluded from the other two gene sets.

Analysis of genes classified as ‘ERα targets’ revealed that ERα loss had no measurable effect on their hypoxia-driven expression, with no significant change in absolute LFC in response to hypoxia following ERα degradation (**Fig. 4b**). Further investigation revealed that this lack of change is driven by the fact the majority of these transcripts do not respond to hypoxia (LFC ≈ 0) in either condition **(Supplementary Fig. S6)**.

By contrast, within the ‘Hypoxia’ gene set, ERα loss significantly blunted the transcriptional response to hypoxia, as evidenced by a reduction in transcript LFCs (p = 9.36 × 10^-183^, paired t-test; **Fig. 4b**). This attenuation was even more pronounced in the ‘Shared Response’ gene set, where ERα depletion reduced the maximum absolute LFC and led to greater reduction in LFCs of DEGs with increased statistical significance (p < 1.0 × 10^-300^, paired t-test; **Fig. 4b**). These findings suggest that ERα activity amplifies the hypoxic transcriptional programme, with the most substantial effect observed among genes co-regulated by both pathways.

To quantify the influence of ERα on the hypoxic transcriptional response, we performed correlation analyses comparing LFCs of transcripts in response to hypoxia in the ERα-expressing and ERα-depleted conditions. A reduced correlation would indicate that ERα loss disrupts the hypoxia-induced transcriptional programme.

‘Hypoxia’ transcripts showed strong linear correlation in the hypoxic response between the two conditions (R = 0.75, p < 0.001, Fisher’s Z test; **Fig. 4c**), indicating the direction and coordination of gene regulation was largely preserved. However, the slope of the linear fit was less than one (0.50), reflecting a general dampening of response magnitude. This result is consistent with **Fig. 4b**, and suggests that while the hypoxic response remains coordinated in the absence of ERα, its intensity is reduced even within our hypoxia specific gene set.

In contrast, our ‘ERα targets’ displayed weak correlation between conditions (R = 0.049, p < 0.001, Fisher’s Z test; **Fig. 4d**), consistent with their insensitivity to hypoxia.

As with our LFC analysis, the ‘Shared Response’ gene set showed the most pronounced ERα-dependent change. Correlation between conditions was reduced relative to the ‘Hypoxia’ gene set (R = 0.58, p < 0.001; **Fig. 4e**), and the slope of the linear fit was further decreased (0.37), indicating a greater dependency on ERα for both amplitude and concordance of the hypoxic transcriptional response.

To control for background expression noise, we analysed genes not included in any of the three defined sets; these transcripts showed only weak correlation between conditions (R = 0.12, p < 0.001, Fisher’s Z test; **Fig. 4f**), as expected.

Together, these results highlight a role for ERα in regulating the hypoxic response in MCF7 cells. Moreover, the extent to which the hypoxic response depends on ERα correlates with how strongly ERα regulates gene expression under normal oxygen conditions (normoxia); i.e, transcripts that have a significant response to changes in ERα activity in normoxia tend to show a greater ERα-dependent modulation during hypoxia.

### ERα Dependence Defines a Novel Hypoxia-Induced Gene Set including ENaC Subunits as Transcriptional Targets

Having established that ERα loss attenuates the hypoxic transcriptional response, we next investigated whether a subset of genes in ER+ breast cancer cells exhibit a dependence on ERα for their hypoxic expression. We defined ERα-dependent hypoxia-responsive transcripts across two cell lines using the following criteria (**Fig. 5a**): (i) significant induction under hypoxia in vehicle-treated cells (adjusted p < 0.01, LFC > 1.5); (ii) significant suppression of hypoxic expression when ERα is degraded with fulvestrant (adjusted p < 0.01, LFC < −1.5; **Supplementary Tables S8-9**); and (iii) no residual induction in response to hypoxia in fulvestrant-treated cells (LFC < 1.5). The final criterion ensures ERα dependence by filtering out transcripts that, while showing a significantly weaker response, still retain partial hypoxic induction in the absence of ERα.

**Figure 5.**
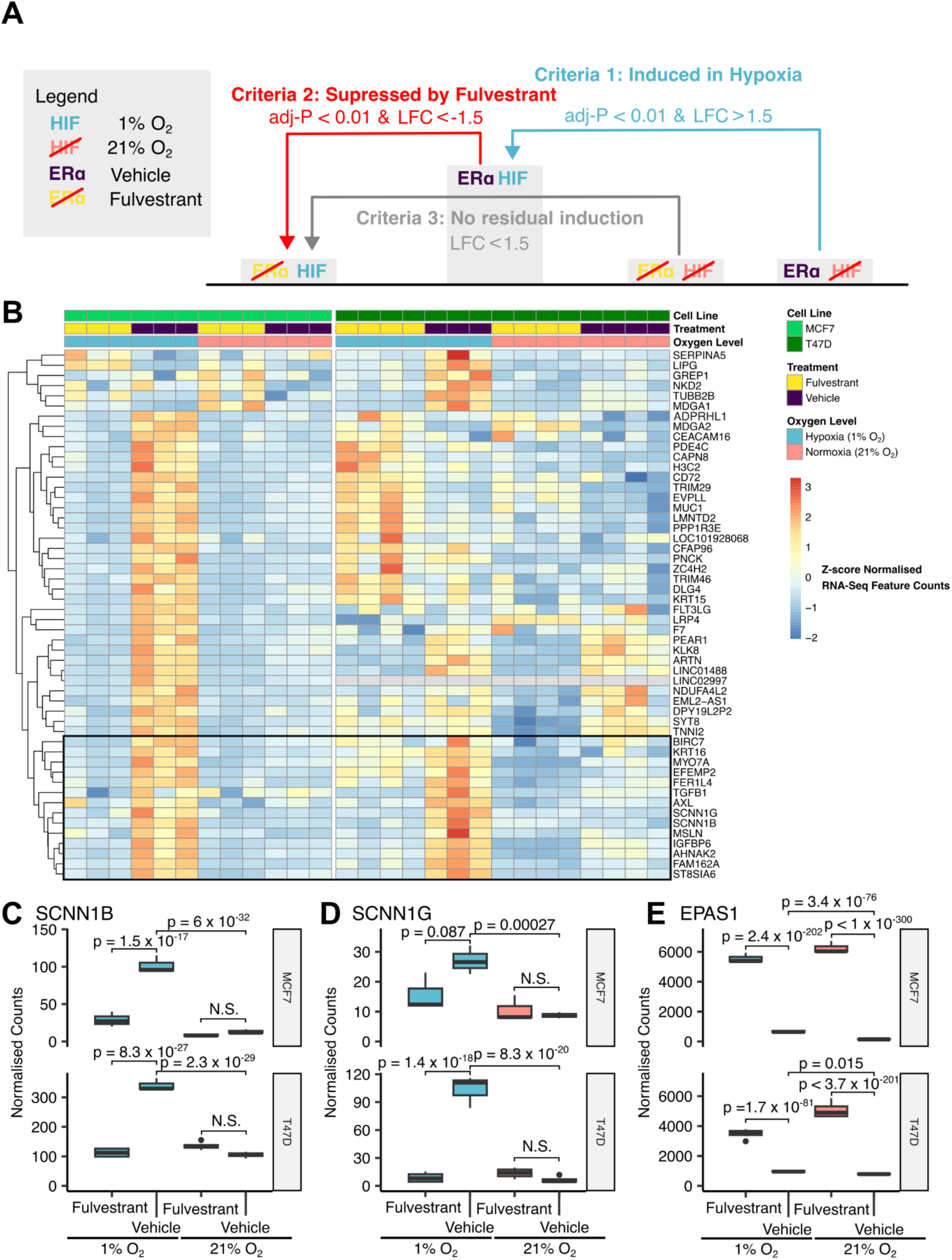
RNA-seq analysis of hypoxia response with and without ERα protein identifies a module of transcripts with an ERα-dependent hypoxia response. (A) Schematic of EDHR selection criteria. (B) DEGs from MCF7 and T47D were selected on the basis that they showed ERα-dependent hypoxic response. Hierarchical clustering of these transcripts identified 14 transcripts that have an EDHR response in both cell lines, including the (C) *SCNN1B* and (D) *SCNN1G* regulatory subunits of ENaC. (E) *EPAS1* (HIF-2α) expression was increased in response to ERα degradation in both MCF7 (p < 1 × 10^-300^) and T47D (p = 3.7 × 10^-201^) without hypoxia.

Across both cell lines, we identified 52 DEGs meeting these criteria (**Fig. 5b**). The majority of these were specific to the MCF7 cell line, with only six unique to the T47D cell line. Hierarchical clustering revealed 14 genes with a shared ERα-dependent hypoxia response (EDHR) across both cell lines.

Among the consensus EDHR genes, several were associated with known EMT, chemoresistance and hypoxia pathways. For example, *TGFB1* is a classic EMT driver in breast cancer (Janus *et al*, 2024; Wang *et al*, 2019), and its transcription and secretion are enhanced under hypoxia (Hung *et al*, 2013)*. AXL,* a HIF1α-associated receptor tyrosine kinase in breast cancer (Nalwoga *et al*, 2016), is typically linked to triple-negative breast cancer and chemoresistance, although its role in ER+ cells has been largely overlooked due to its lack of expression under normoxic conditions (Meyer *et al*, 2013). *KRT16* has been associated with EMT and metastatic progression (Joosse *et al*, 2012; Elazezy *et al*, 2021), while *BIRC5* (encoding survivin) is associated with resistance to tamoxifen-induced apoptosis (Moriai *et al*, 2009).

In addition, we observed selective, ERα-dependent hypoxic induction of *SCNN1B* (**Fig. 5c**) and *SCNN1G* (**Fig. 5d**), which encode the β and γ subunits of the epithelial sodium channel (ENaC), respectively, in both MCF7 and T47D cells.

As a positive control, we confirmed that the loss of ERα induced a significant increase in *EPAS1* transcript levels both under hypoxia and normoxia (**Fig. 5e**). This observation is consistent with previous reports describing estrogen-mediated repression of HIF-2α (Fuady *et al*, 2016), and confirms that this regulatory interaction is active in our culture conditions.

These findings extend previous observations of hormone-hypoxia crosstalk and suggest a novel regulatory axis involving ENaC subunits downstream of a ERα-HIF interaction.

### ENaC Regulatory Subunits Predict ERα⁺ Breast Cancer Outcomes and Show Hypoxia-Dependent Drug Sensitivity

On the basis that the genes encoding β-ENaC and γ-ENaC were both selectively responsive to ERα-dependent hypoxic induction, we aimed to investigate the potential relevance of these subunits in the context of ER+ breast cancer. ENaC is a heterotrimer composed of a Na^+^-transporting α subunit, encoded by *SCNN1A*, and the two regulatory β and γ subunits (Hamm *et al*, 2010). Various Na^+^ channels, including ENaC, are aberrantly expressed in cancer cells (Leslie *et al*, 2019; Malcolm *et al*, 2023), and altered Na^+^ transport has previously been shown to have significant effects on the hypoxia response by preventing mitochondrial adaptation to hypoxia (Hernansanz-Agustín *et al*, 2020). Furthermore, the ENaC inhibitor, amiloride, has been shown to increase viability in normoxia for MCF7 and T47D cells (Ware *et al*, 2021). Based on these findings, we investigated whether the expression of the ENaC regulatory subunits was associated with breast cancer survival, and whether hypoxia modulates cellular responses to amiloride.

We initially utilised the Kaplan-Meier Plotter (Győrffy, 2024) to stratify ER+ breast cancer patients into high and low expression groups of *SCNN1B* and *SCNN1G,* and to predict RFS. Low expression of *SCNN1B* (**Fig. 6a**) and *SCNN1G* (**Fig. 6b**) was significantly associated with better RFS (p = 0.015, p = 2.5 x 10^-6^, respectively), whereas high expression of these subunits was prognostic of poorer outcomes in ER+ breast cancer. Interestingly, this trend contrasts with that reported for *SCNN1A*, where high expression is associated with improved overall survival when not stratified by subtype (OS, p = 0.002) (Ware *et al*, 2021). We reproduced this result, showing a similar association with OS (p = 0.044) and a stronger association with RFS (p = 8 x 10^-12^).

**Figure 6.**
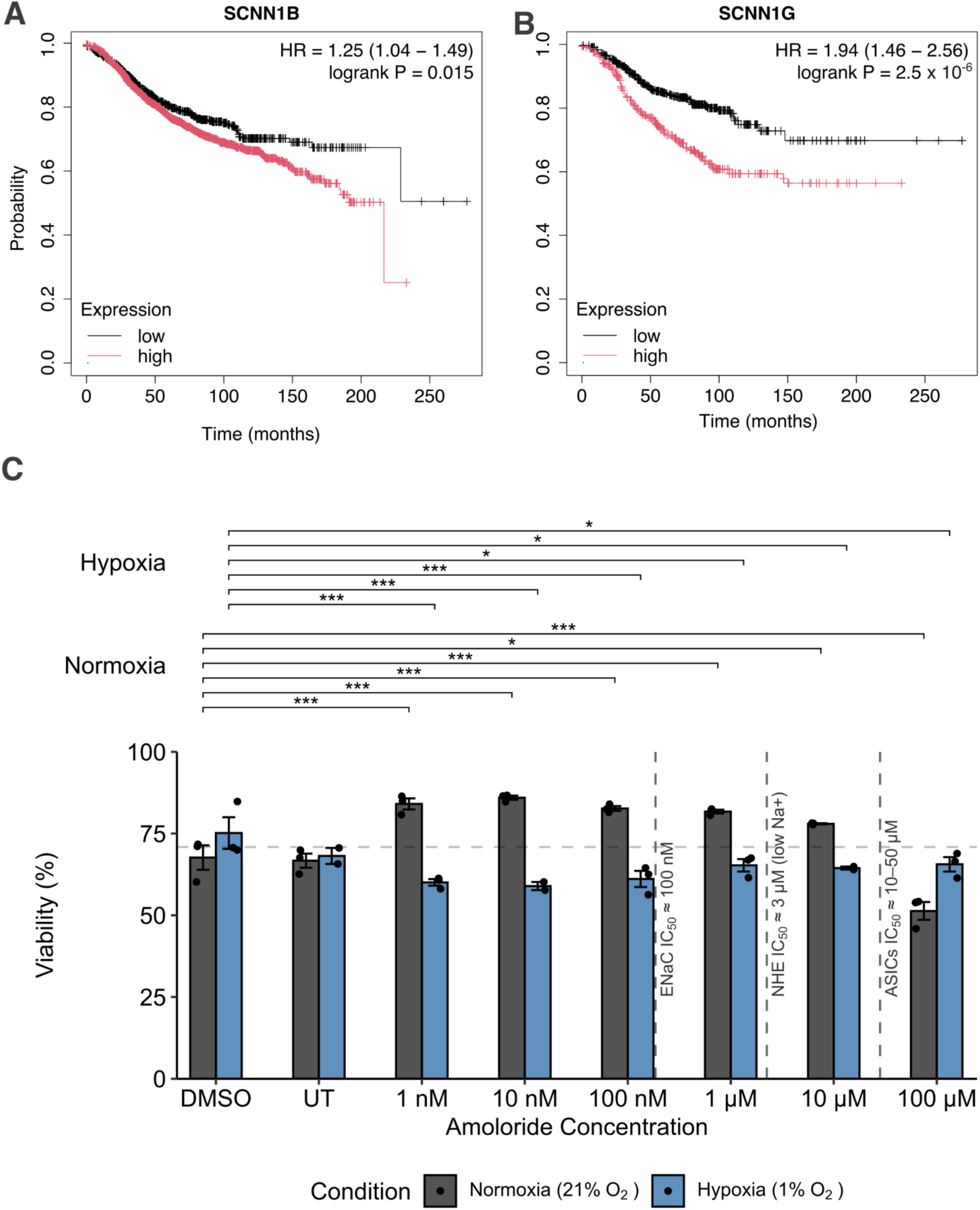
High expression of ENaC regulatory subunits is implicated with shorter RFS, and the ENaC inhibitor, amiloride, alters viability in hypoxic MCF7 cells. RFS analysis of ENaC subunits *SCNN1B* (A) and *SCNN1G* (B) demonstrate a significant association with low expression of these ENAC regulatory subunits and better patient outcome. (C) Dose-response analysis of amiloride-treated MCF7 cells under normoxia and hypoxia reveals hypoxia-specific sensitivity. UT = untreated control. Dashed vertical lines indicate approximate IC₅₀ values for ENaC (100 nM), NHE1 (3 μM, low [Na^+^]), and ASICs (10–50 μM). Error bars indicate SEM. Significance: * = p < 0.05, ** = p < 0.01, *** = p < 0.001, ns = not significant.

To further explore this potential contradiction, we stratified by ERα status. *SCNN1A* was not significantly associated with RFS (p=0.068, with low expression trending to worse outcome) or OS (p = 0.23, where high expression was trending to worse outcome); however, in ERα-disease, high expression was associated with worse RFS (p = 0.01) but not OS (p = 0.14; **Supplementary Fig. S7**).

Amiloride is a pleiotropic inhibitor of multiple ion channels and transporters, including the ENaC (0.1 µM), ASICs (10-50 µM), NHE (3 μM to 1 mM depending on Na⁺ conditions) and NCX (1 mM) (Qadri *et al*, 2012; Xiong *et al*, 2008; Teiwes & Toto, 2007). While inhibition of the ENaC has been linked to increased cell viability, higher concentrations of amiloride exhibit cytotoxic effects, particularly through inhibition of NHE1 (IC_50_ = 100 µM (Hu *et al*, 2024)). To address these concerns, we tested a concentration range spanning the electrophysiologically determined amiloride IC₅₀ for ENaC (∼100 nM) (Noreng *et al*, 2020).

To determine whether ENaC inhibition modulates cell viability under hypoxia, MCF7 cells were treated with amiloride (0–100 μM), followed by 72 hours culture in either normoxia or hypoxia (1% oxygen). Viability was assessed by flow cytometry using live/dead staining (**Fig. 6C**).

A two-way ANOVA model with amiloride treatment, oxygen condition, and their interaction as factors revealed significant effects for both treatment (p = 1.12 × 10⁻⁶) and oxygen condition (p = 3.18 × 10⁻⁹), as well as a significant interaction (p = 2.67 × 10⁻¹⁰), indicating that the effect of amiloride on cell viability is dependent on oxygen availability.

Post-hoc pairwise comparisons (Sidak-adjusted, referenced to DMSO vehicle control) showed that under normoxia, amiloride significantly increased viability at concentrations from 1 nM to 10 μM, while 100 μM led to a significant decrease, consistent with cytotoxic effects attributed to NHE inhibition in sodium containing culture medium. In contrast, hypoxic cells showed a significantly reduced viability compared to DMSO control at all amiloride concentrations from 1 nM to 100 μM. No significant differences were detected between DMSO control and the untreated control.

These findings support a model in which ERα-dependent induction of ENaC regulatory subunits enhances hypoxia tolerance, and highlight the context-dependent effects of amiloride in breast cancer cells. Together, they suggest ENaC may represent a hypoxia-responsive vulnerability that could be exploited therapeutically in ER+ tumours.

### The ERα-Dependent Hypoxic Signature is Spatially Associated with EMT in ER+ Tumours

To investigate the intratumoral expression of our consensus-derived 14 gene ERα-dependent hypoxic response, we interrogated spatial transcriptomic profiles of three ER+ breast tumours from publicly available datasets (Withnell & Secrier, 2024). Spatial mapping revealed colocalisation of the MSigDB Estrogen Response Late and Hypoxia hallmark signatures throughout tumour sections, with our EDHR signature frequently bordering these regions and with adjacent areas enriched for EMT (**Fig. 7a**). Given the spatial proximity of our EDHR signature to EMT-enriched regions, we next asked whether our novel signature had a quantifiable association with these areas. Using *SpottedPy (Withnell & Secrier, 2024),* we measured the spatial transcriptomic associations and found a significant spatial association of EMT with EDHR related transcripts, and that EDHR was the closest of the three studied signatures to sites of EMT (**Fig. 7b**). Moreover, correlation analysis with *SpottedPy* between signatures in the neighbourhood of each tumour spot similarly showed a correlation between our EDHR signature and EMT (**Fig. 7c**).

**Figure 7.**
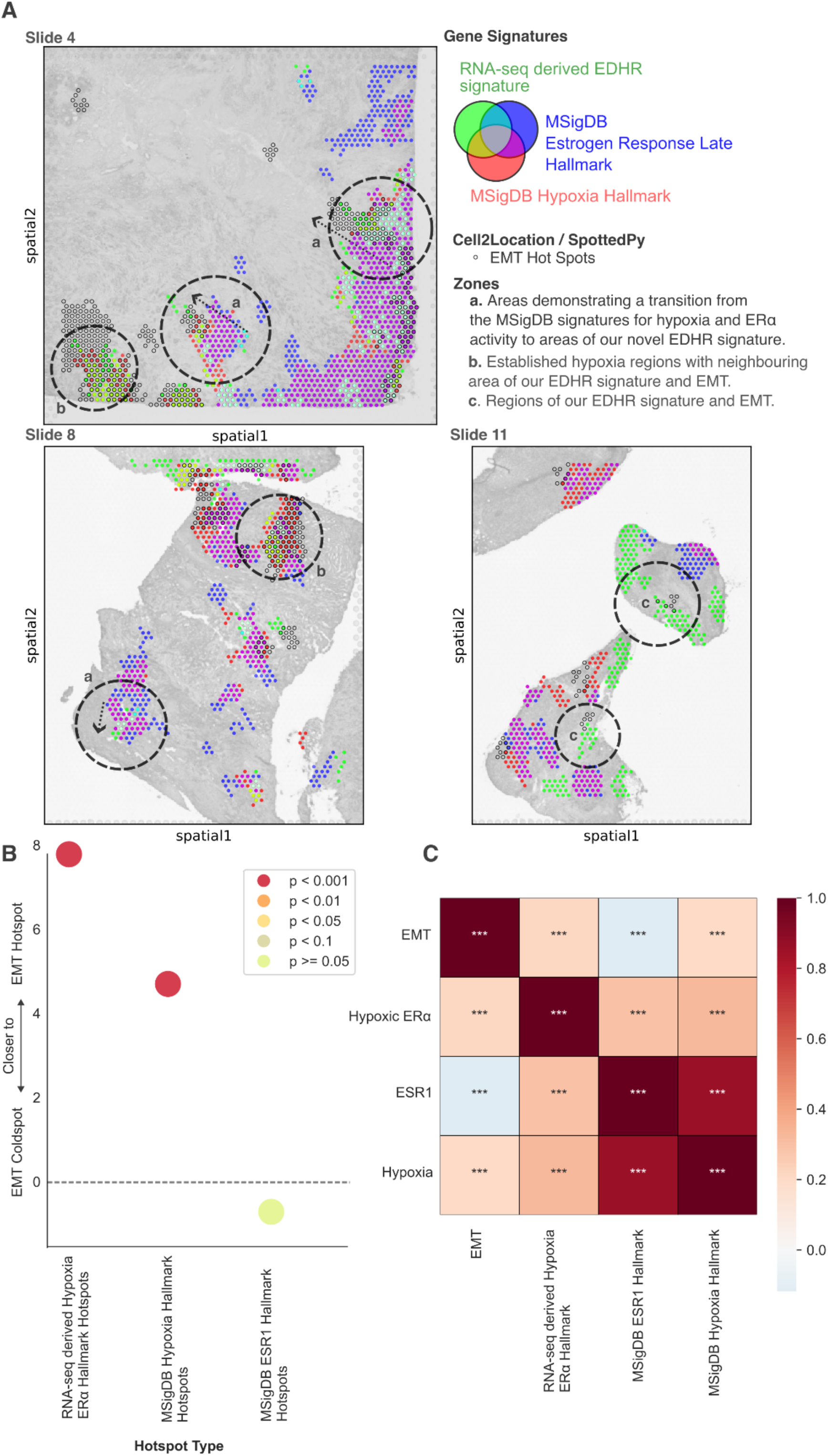
Analysis of spatial transcription data of ER+ patient tumours shows a significant spatial association of our EDHR and EMT. (A) Spatial analysis of ERα tumours showed that the MSigDB hallmarks for *Estrogen Response Late* and *Hypoxia* coexisted through the tumour, with our EDHR transcripts often on the peripheral of these regions and close to regions of EMT. (B) Analysis of the visual data in (A) using *SpottedPy* to quantify these data confirmed a significant spatial association of EMT with our EDHR (p < 0.001), and EDHR was the closest signature to sites of EMT. (C) Correlation analysis by *SpottedPy* between signatures in the neighbourhood of each tumour spot similarly showed a correlation between our EDHR signature and EMT.

Together, these findings indicate that the ERα-dependent hypoxic programme is not only spatially organised within ERα⁺ tumours but is also preferentially enriched near EMT regions, and significantly associated with areas of hypoxia.

## Discussion

In this study, we describe the genome-wide redistribution of ERα binding in response to the hypoxia mimetic and dioxygenase inhibitor DMOG. We also observe widescale ERα redistribution to new binding sites after 16 hours of treatment with DMOG. Analysis of published ATAC-seq data (Miar *et al*, 2020) indicates that these ERα binding events occur at regions of pre-existing chromatin accessibility, with no detectable changes after 48 hours at 0.1% oxygen. This finding suggests that chromatin opening is not the primary driver of ERα reprogramming under these conditions. Given the oxygen dependence of histone demethylases, and their inhibition by DMOG, a histone methylation status may well be a contributor alongside the activation of HIFs. Indeed, increases in histone methylation have previously been reported under hypoxic conditions in MCF7 cells exposed to hypoxia (Prickaerts *et al*, 2016). Although histone methylation does not affect histone charge, so does not directly alter chromatin compaction, it can influence transcriptional outcomes by regulating the recruitment of chromatin-associated effector complexes (Bannister & Kouzarides, 2011).

Strikingly, ERα binding alterations under DMOG treatment were enriched for FOXA1 motifs as well as EREs, suggesting that ERE binding by ERα remains relevant. This implies that epigenomic modifications or altered cofactor availability may modulate ERα affinity at these loci. Interactions with other transcriptional regulators such as HIF-1β (ARNT) may also contribute, given that it can modulate ERα chromatin binding (Madak-Erdogan & Katzenellenbogen, 2012).

Whether the redistribution of ERα is primarily driven by HIF stabilisation, epigenomic changes, or both remains unresolved. However, the use of more selective PHD inhibitors, such as IOX5 (Lawson *et al*, 2024), which selectively activates HIF-1α without broadly inhibiting demethylases, may help disentangle these effects.

Integration of our ChIP-seq and transcriptomic data generated two ERα-dependent signatures defined by DMOG-induced redistribution. PCA of 31 different cell lines (Godet *et al*, 2019) based on our identified gene signatures, showed that ER+ cell lines specifically clustered together for both loss- and gain-associated signatures (**Fig. 2a,b**). We proceeded to show that our gene signatures are prognostically relevant, as patients with high expression of loss-associated genes show greater RFS; and the opposing prognostic association was found for gain-associated gene expression (**Fig. 2d,e**) (Cancer Genome Atlas Network, 2012). Through the inclusion of several breast cancer subtypes in this analysis, we can confirm that these hypoxia-induced alterations are specific to ERα-positive cell lines. Further, given that the unique set of genes does not overlap with a highly prognostic metagene (Buffa *et al*, 2010), we can also be confident that these changes are not due to the generalised hypoxic response.

Most previous studies examining ERα-HIF crosstalk have focused on either hierarchical regulation of transcription factor expression or functional compensation between the pathways. For example, HIF-1α activity has been shown to downregulate ERα expression (Wolff *et al*, 2017; Padró *et al*, 2017), while ERα can enhance *HIF1A* transcription and repress *EPAS1* via EREs (Yang *et al*, 2015; Fuady *et al*, 2016). In addition, estrogen signalling also modulates the HIF expression through transcriptional upregulation of PHD1 (Appelhoff *et al*, 2004). Compensatory mechanisms include the ability of ERα and HIF-1α to interchangeably drive VEGFA expression and modulate epithelial-endothelial interactions (George *et al*, 2012), HIF-1α compensation for loss of ERα in the regulation of SNAT2 (Morotti *et al*, 2019), and the co-regulation of JMJD2B (Yang *et al*, 2010).

These studies therefore demonstrate a bidirectional interaction between ERα and HIFs, yet they largely reflect parallel or upstream regulatory effects and present limited insight into the potential for integration of the two signalling pathways. In contrast, our observation of widespread ERα redistribution under hypoxia-like conditions suggests a functional role for ERα in shaping the transcriptional hypoxic response in ER+ cells.

To the best of our knowledge, only one other study has previously reported widespread remodelling of the ERα cistrome in MCF7 cells under hypoxia (Jehanno *et al*, 2022). However, the experimental design within that study asks an important, but different, question; they examined how long-term hypoxia (1 month of culture in the presence of hypoxia mimetic CoCl₂) shapes the response by ERα. In that context, ERα binding was largely lost, and gene regulation was maintained through HIF-driven enhancer activity with only a small number of hypoxia-specific ERα binding sites. Thus, the dependence of the hypoxic response on ERα in ER+ cells has not been previously characterised.

To address this gap, we performed RNA-seq measuring the response to hypoxia (1% oxygen, 48 hours) and ERα degradation (via fulvestrant, 96 hours). Differential gene expression analysis reveals that 41% of DEGs identified were shared across both conditions (**Fig. 4**). However, this overlap alone does not distinguish between convergent pathway effects and functional interdependence. We therefore performed a combined treatment, using fulvestrant to ablate ERα before culturing in hypoxia, in order to define an ERα-Dependent Hypoxia Response (EDHR).

In response to ERα loss, we observed a marked reduction in the transcription of hypoxia-responsive genes, indicating a strong functional relationship between ERα and hypoxic signalling in ER+ cells. Correlation analysis (**Fig. 4c-f**) of our hypoxia-specific DEGs confirmed strong agreement with the response seen in our single-factor experiments, albeit with reduced Log_2_ Fold Changes (LFCs). In contrast, DEGs within the shared gene set (i.e. both ERα- and hypoxia-regulated) exhibited significantly reduced correlation and attenuated expression, consistent with dual regulation by both factors.

We interpret these findings to suggest that as loss of ERα leads to decreased HIF1α levels (Yang *et al*, 2015), there is a dampened hypoxic transcriptional response across all targets. However, for those with a shared response, this effect is compounded by the absence of ERα regulation, producing greater attenuation of expression and reduced correlation. This model aligns with the known dual role of ERα in both activating and repressing hypoxia-related genes, as demonstrated by its positive regulation of *HIF1A* and repression of *EPAS1* through EREs (**Fig. 3c**) (Fuady *et al*, 2016; Yang *et al*, 2015).

Across both MCF7 and T47D cells, we identified 14 EDHR genes that were ERα-dependent under hypoxia, with minimal detectable expression in normoxia or following ERα degradation by fulvestrant (**Fig. 5**). Notably, we identified two regulatory subunits of the protein ENaC: β-ENaC and γ-ENaC as EDHR targets. Interestingly, the γ-ENaC subunit has been linked to pro-carcinogenic effects in high-salt microenvironments (Amara *et al*, 2016). In contrast, the pore-forming α-ENaC subunit has recently been shown to slow the proliferation and migration of breast cancer cells (McQueen *et al*, 2025; Ware *et al*, 2021).

Expression of the ENaC sodium channel has also been linked to p53 mutation status, ER status, and histological tumour grade (Ko *et al*, 2013). Although best characterised for their canonical role in renal sodium reabsorption and systemic blood pressure regulation (Shimkets *et al*, 1994; Strautnieks *et al*, 1996; Canessa *et al*, 1994; Oxlund *et al*, 2014; Kremer *et al*, 1977), ENaC subunits have emerging roles in non-renal tissues, including in mechanotransduction pathways that promote cell migration and invasiveness (Xu *et al*, 2012; Grifoni *et al*, 2006; Yang *et al*, 2013; Jeggle *et al*, 2013; Drummond *et al*, 2008). Consistent with an oncogenic role, analysis of RFS in breast cancer patients revealed that high expression of *SCNN1B* and *SCNN1G* was significantly associated with poorer prognosis (**Fig. 6**), suggesting that EDHR-mediated upregulation of these subunits may be clinically relevant in ER+ disease.

To investigate the functional relevance of EDHR-induced ENaC expression, we tested the effect of the ENaC inhibitor amiloride on the viability of normoxic and hypoxic MCF7 cells. Under normoxia, amiloride enhanced viability at lower concentrations (1 nM–10 μM) but became cytotoxic at 100 μM, likely due to NHE1 inhibition (Hu *et al*, 2024). Notably, under hypoxia, amiloride had the opposite effect, reducing viability at all tested concentrations. These data suggest that ERα-dependent induction of ENaC subunits by the EDHR may support hypoxia tolerance in ER+ cells. These findings reveal a potential context-specific vulnerability in hormone-driven breast cancer.

Finally, to assess the clinical relevance of the ERα-dependent hypoxic response *in situ*, we examined its spatial organisation in ERα⁺ breast tumours using publicly available spatial transcriptomics data sets. Across all tumours, EDHR expression was detected, with spatial mapping revealing colocalisation of the MSigDB ‘Estrogen Response Late’ and ‘Hypoxia’ hallmark gene signatures throughout tumour sections, while the EDHR signature was enriched at their margins and frequently adjacent to regions expressing EMT markers (**Fig. 7a**).

Quantitative analysis with *SpottedPy* revealed that EMT-related transcripts were significantly associated with EDHR-expressing regions, and that EDHR was consistently the closest of the three signatures to EMT hotspots (**Fig. 7b**). Correlation analysis across spatial neighbourhoods further confirmed a significant association between localisation of the EDHR and EMT signatures.

A limitation of our study is the inability to definitively distinguish whether the observed EDHR gene expression arises from direct ERα binding at target loci, or instead reflects secondary effects involving additional transcription factor activity or broader epigenetic changes. This complexity is particularly relevant given the extended (96-hour) fulvestrant treatment used to achieve complete ERα degradation. However, fulvestrant is a well-characterised selective estrogen receptor degrader (SERD), and we are confident in its specificity and effectiveness. Thus, while the precise mechanism of regulation may vary, our data robustly support a requirement for ERα in the hypoxic induction of these transcripts in ER+ breast cancer.

Although the number of defined EDHR genes was modest (n = 14), our transcriptome-wide correlation analysis indicates that this represents a subset of a broader spectrum of ERα dependency. Less stringent thresholds may capture additional genes with partial dependency or alternative models of integration with ERα signalling, providing a basis for future mechanistic studies.

In conclusion, we identify a set of hypoxia-responsive genes with dependence on ERα in ER+ breast cancer, revealing novel mechanisms of ERα signalling. While previous studies have primarily focused on hypoxia-induced epigenetic remodelling or transcriptional resistance mechanisms, our findings demonstrate that ERα itself is a key coordinator of a distinct hypoxic gene programme in ER+ breast cancer. Echoing recent reports regarding NF-κB as a central regulator of hypoxia-induced gene expression (Shakir *et al*, 2025).

We highlight the relevance of the ENaC subunits encoded by *SCNN1B* and *SCNN1G* as EDHR targets, demonstrating an association with poor prognosis, and we show that pharmacological inhibition of ENaC selectively impairs hypoxic viability, suggesting a novel therapeutic vulnerability.

Further, spatial transcriptomic analysis of ER+ tumours revealed consistent localisation of the EDHR signature adjacent to EMT-enriched regions, highlighting not only the clinical presence of this transcriptional response, but its spatial association with aggressive tumour features. These results provide strong evidence that the EDHR contributes to tumour progression *in vivo*.

Taken together, our study identifies a role for ERα in shaping the hypoxic response in ER+ breast cancer, with translational relevance for both prognosis and therapeutic intervention.

## Data availability

Custom scripts and notebooks used for statistical analysis and figure generation are available at https://github.com/Holding-Lab/ER-HIF (Holding, 2025), and undertaken using R (v4.4.0) or Python (v3.2.17). Differential expression was undertaken using DESeq2 (Love *et al*, 2014), and figures were generated using ggplot2 (Wickham, 2016).

The raw ChIP-seq data generated in this study have been deposited in ArrayExpress under accession number E-MTAB-15329. The raw RNA-seq data are available under accession number E-MTAB-15331.

## Author contributions

JRM and JS contributed equally to this work. As per CRediT taxonomy. Conceptualization, KSB, WJB, ANH; Methodology, JRM, JP, JŁ, SFR, TES; Software, ANH; Validation, TES, JŁ, JP; Formal Analysis, JRM, JŁ, TES, ANH; Investigation, JRM, JP, JŁ, SFR, TES; Resources, LG, SRJ; Data Curation LG, SRJ, ANH; Visualization, ANH, JP, JRM; Writing (Original Draft), JRM, JS, ANH; Writing (Review & Editing), all authors; Supervision, KSB, WJB, ANH; Project Administration, ANH; Funding Acquisition, KSB, WJB, ANH.

## Acknowledgements

This work was funded by the Biotechnology and Biological Sciences Research Council (BBSRC) (BB/V000071/1 & BB/X511213/1) to ANH, and (BB/T007222/1) as PhD studentships to JRM and JS. WJB reports funding from York Against Cancer, MRC (MR/X018067/1), BBSRC (BB/Y513970/1), and the Wellcome Trust (310891/Z/24/Z).The authors would like to thank the Bioscience Technology Facility, University of York for their support in Genomic workflows and Flow Cytometry analysis. We are grateful to Professor David Kent (University of York) for facilitating support towards RNA-seq experiments. The Viking cluster was used during this project, which is a high-performance compute facility provided by the University of York. We are grateful for computational support from the University of York, IT Services and the Research IT team. The results published here are in part based upon data generated by the TCGA Research Network: https://www.cancer.gov/tcga.

## Abbreviations

ATAC-Seq: Assay for Transposase-Accessible Chromatin with Sequencing
BP: Biological Process
ChIP-Seq: Chromatin Immunoprecipitation with Sequencing
CoCl_2_: Cobalt Chloride
DEGs: Differentially Expressed Genes
DMOG: Dimethyloxalylglycine
ECM: Extracellular Matrix
EDHR: ERα-Dependent Hypoxic Response
EMT: Epithelial-to-Mesenchymal Transition
ENaC: Epithelial Sodium Channel
ERα: Estrogen Receptor α
ERE: Estrogen Response Element
FAD: Flavin Adenine Dinucleotide
GO: Gene Ontology
HIF: Hypoxia Inducible Factor
HRE: Hypoxia Response Element
LFC: Log Fold Change
O_2_: Oxygen
RFS: Recurrence-Free Survival
TNBC: Triple Negative Breast Cancer
HMA: Hypoxia Mimetic Agent

## Notes

https://doi.org/10.5281/zenodo.17221105

### Competing Interest Statement

The authors have declared no competing interest.

## References

Amara S, Ivy MT, Myles EL & Tiriveedhi V (2016) Sodium channel γENaC mediates IL-17 synergized high salt induced inflammatory stress in breast cancer cells. Cell Immunol 302: 1–10

Amemiya HM, Kundaje A & Boyle AP (2019) The ENCODE blacklist: Identification of problematic regions of the genome. Sci Rep 9: 9354

Appelhoff RJ, Tian Y-M, Raval RR, Turley H, Harris AL, Pugh CW, Ratcliffe PJ & Gleadle JM (2004) Differential function of the prolyl hydroxylases PHD1, PHD2, and PHD3 in the regulation of hypoxia-inducible factor. J Biol Chem 279: 38458–38465

Bannister AJ & Kouzarides T (2011) Regulation of chromatin by histone modifications. Cell Res 21: 381–395

Baron S, Escande A, Albérola G, Bystricky K, Balaguer P & Richard-Foy H (2007) Estrogen receptor alpha and the activating protein-1 complex cooperate during insulin-like growth factor-I-induced transcriptional activation of the pS2/TFF1 gene. J Biol Chem 282: 11732–11741

Batie M, Frost J, Shakir D & Rocha S (2022) Regulation of chromatin accessibility by hypoxia and HIF. Biochem J 479: 767–786

Bracken CP, Fedele AO, Linke S, Balrak W, Lisy K, Whitelaw ML & Peet DJ (2006) Cell-specific regulation of hypoxia-inducible factor (HIF)-1alpha and HIF-2alpha stabilization and transactivation in a graded oxygen environment. J Biol Chem 281: 22575–22585

Brunnberg S, Pettersson K, Rydin E, Matthews J, Hanberg A & Pongratz I (2003) The basic helix-loop-helix-PAS protein ARNT functions as a potent coactivator of estrogen receptor-dependent transcription. Proc Natl Acad Sci U S A 100: 6517–6522

Buffa FM, Harris AL, West CM & Miller CJ (2010) Large meta-analysis of multiple cancers reveals a common, compact and highly prognostic hypoxia metagene. Br J Cancer 102: 428–435

Cancer Genome Atlas Network (2012) Comprehensive molecular portraits of human breast tumours. Nature 490: 61–70

Canessa CM, Schild L, Buell G, Thorens B, Gautschi I, Horisberger JD & Rossier BC (1994) Amiloride-sensitive epithelial Na+ channel is made of three homologous subunits. Nature 367: 463–467

Capatina AL, Malcolm JR, Stenning J, Moore RL, Bridge KS, Brackenbury WJ & Holding AN (2024) Hypoxia-induced epigenetic regulation of breast cancer progression and the tumour microenvironment. Front Cell Dev Biol 12: 1421629

Carroll JS, Liu XS, Brodsky AS, Li W, Meyer CA, Szary AJ, Eeckhoute J, Shao W, Hestermann EV, Geistlinger TR, et al (2005) Chromosome-wide mapping of estrogen receptor binding reveals long-range regulation requiring the forkhead protein FoxA1. Cell 122: 33–43

Casillas AL, Chauhan SS, Toth RK, Sainz AG, Clements AN, Jensen CC, Langlais PR, Miranti CK, Cress AE & Warfel NA (2021) Direct phosphorylation and stabilization of HIF-1α by PIM1 kinase drives angiogenesis in solid tumors. Oncogene 40: 5142–5152

Chan MC, Ilott NE, Schödel J, Sims D, Tumber A, Lippl K, Mole DR, Pugh CW, Ratcliffe PJ, Ponting CP, et al (2016) Tuning the transcriptional response to hypoxia by inhibiting hypoxia-inducible factor (HIF) prolyl and asparaginyl hydroxylases. J Biol Chem 291: 20661–20673

Chen C, Pore N, Behrooz A, Ismail-Beigi F & Maity A (2001) Regulation of glut1 mRNA by hypoxia-inducible factor-1. Interaction between H-ras and hypoxia. J Biol Chem 276: 9519–9525

Chen R, Ahmed MA & Forsyth NR (2022) Dimethyloxalylglycine (DMOG), a hypoxia mimetic agent, does not replicate a rat pheochromocytoma (PC12) cell biological response to reduced oxygen culture. Biomolecules 12: 541

Collier H, Albanese A, Kwok C-S, Kou J & Rocha S (2023) Functional crosstalk between chromatin and hypoxia signalling. Cell Signal 106: 110660

Culhane JC & Cole PA (2007) LSD1 and the chemistry of histone demethylation. Curr Opin Chem Biol 11: 561–568

Drummond HA, Grifoni SC & Jernigan NL (2008) A new trick for an old dogma: ENaC proteins as mechanotransducers in vascular smooth muscle. Physiology (Bethesda) 23: 23–31

Duan C (2016) Hypoxia-inducible factor 3 biology: complexities and emerging themes. Am J Physiol Cell Physiol 310: C260–9

Elazezy M, Schwentesius S, Stegat L, Wikman H, Werner S, Mansour WY, Failla AV, Peine S, Müller V, Thiery JP, et al (2021) Emerging insights into keratin 16 expression during metastatic progression of breast cancer. Cancers (Basel) 13: 3869

Figg WD Jr, McDonough MA, Chowdhury R, Nakashima Y, Zhang Z, Holt-Martyn JP, Krajnc A & Schofield CJ (2021) Structural basis of prolyl hydroxylase domain inhibition by Molidustat. ChemMedChem 16: 2082–2088

Forsythe JA, Jiang BH, Iyer NV, Agani F, Leung SW, Koos RD & Semenza GL (1996) Activation of vascular endothelial growth factor gene transcription by hypoxia-inducible factor 1. Mol Cell Biol 16: 4604–4613

Fuady JH, Gutsche K, Santambrogio S, Varga Z, Hoogewijs D & Wenger RH (2016) Estrogen-dependent downregulation of hypoxia-inducible factor (HIF)-2α in invasive breast cancer cells. Oncotarget 7: 31153–31165

Fu X, Pereira R, De Angelis C, Veeraraghavan J, Nanda S, Qin L, Cataldo ML, Sethunath V, Mehravaran S, Gutierrez C, et al (2019) FOXA1 upregulation promotes enhancer and transcriptional reprogramming in endocrine-resistant breast cancer. Proc Natl Acad Sci U S A 116: 26823–26834

Garcia-Bassets I, Kwon Y-S, Telese F, Prefontaine GG, Hutt KR, Cheng CS, Ju B-G, Ohgi KA, Wang J, Escoubet-Lozach L, et al (2007) Histone methylation-dependent mechanisms impose ligand dependency for gene activation by nuclear receptors. Cell 128: 505–518

George AL, Rajoria S, Suriano R, Mittleman A & Tiwari RK (2012) Hypoxia and estrogen are functionally equivalent in breast cancer-endothelial cell interdependence. Mol Cancer 11: 80

Glont S-E, Papachristou EK, Sawle A, Holmes KA, Carroll JS & Siersbaek R (2019) Identification of ChIP-seq and RIME grade antibodies for Estrogen Receptor alpha. PLoS One 14: e0215340

Godet I, Shin YJ, Ju JA, Ye IC, Wang G & Gilkes DM (2019) Fate-mapping post-hypoxic tumor cells reveals a ROS-resistant phenotype that promotes metastasis. Nat Commun 10: 4862

Grifoni SC, Gannon KP, Stec DE & Drummond HA (2006) ENaC proteins contribute to VSMC migration. Am J Physiol Heart Circ Physiol 291: H3076–86

Gu YZ, Moran SM, Hogenesch JB, Wartman L & Bradfield CA (1998) Molecular characterization and chromosomal localization of a third alpha-class hypoxia inducible factor subunit, HIF3alpha. Gene Expr 7: 205–213

Győrffy B (2024) Integrated analysis of public datasets for the discovery and validation of survival-associated genes in solid tumors. Innovation (Camb) 5: 100625

Hamm LL, Feng Z & Hering-Smith KS (2010) Regulation of sodium transport by ENaC in the kidney. Curr Opin Nephrol Hypertens 19: 98–105

Hayashi M, Sakata M, Takeda T, Yamamoto T, Okamoto Y, Sawada K, Kimura A, Minekawa R, Tahara M, Tasaka K, et al (2004) Induction of glucose transporter 1 expression through hypoxia-inducible factor 1alpha under hypoxic conditions in trophoblast-derived cells. J Endocrinol 183: 145–154

Hernansanz-Agustín P, Choya-Foces C, Carregal-Romero S, Ramos E, Oliva T, Villa-Piña T, Moreno L, Izquierdo-Álvarez A, Cabrera-García JD, Cortés A, et al (2020) Na+ controls hypoxic signalling by the mitochondrial respiratory chain. Nature 586: 287–291

Higashimura Y, Nakajima Y, Yamaji R, Harada N, Shibasaki F, Nakano Y & Inui H (2011) Up-regulation of glyceraldehyde-3-phosphate dehydrogenase gene expression by HIF-1 activity depending on Sp1 in hypoxic breast cancer cells. Arch Biochem Biophys 509: 1–8

Hodgkinson K, Forrest LA, Vuong N, Garson K, Djordjevic B & Vanderhyden BC (2018) GREB1 is an estrogen receptor-regulated tumour promoter that is frequently expressed in ovarian cancer. Oncogene 37: 5873–5886

Holding, A. N. (2025). Holding-Lab/ER-HIF: bioRxiv v1 - preprint (preprint). Zenodo. 10.5281/zenodo.17221105

Holding AN, Cullen AE & Markowetz F (2018) Genome-wide Estrogen Receptor-α activation is sustained, not cyclical. Elife 7

Hopkinson RJ, Tumber A, Yapp C, Chowdhury R, Aik W, Che KH, Li XS, Kristensen JBL, King ONF, Chan MC, et al (2013) 5-Carboxy-8-hydroxyquinoline is a Broad Spectrum 2-Oxoglutarate Oxygenase Inhibitor which Causes Iron Translocation. Chem Sci 4: 3110–3117

Huang H, Wei T, Zhang A, Zhang H, Kong L, Li Y & Li F (2024) Trends in the incidence and survival of women with hormone receptor-positive breast cancer from 1990 to 2019: a large population-based analysis. Sci Rep 14: 23690

Hu M, Liu R, Castro N, Loza Sanchez L, Rueankham L, Learn JA, Huang R, Lam KS & Carraway KL 3rd (2024) A novel lipophilic amiloride derivative efficiently kills chemoresistant breast cancer cells. Sci Rep 14: 20263

Hung S-P, Yang M-H, Tseng K-F & Lee OK (2013) Hypoxia-induced secretion of TGF-β1 in mesenchymal stem cell promotes breast cancer cell progression. Cell Transplant 22: 1869–1882

Hurtado A, Holmes KA, Ross-Innes CS, Schmidt D & Carroll JS (2011) FOXA1 is a key determinant of estrogen receptor function and endocrine response. Nat Genet 43: 27–33

Iriyama T, Wang W, Parchim NF, Song A, Blackwell SC, Sibai BM, Kellems RE & Xia Y (2015) Hypoxia-independent upregulation of placental hypoxia inducible factor-1α gene expression contributes to the pathogenesis of preeclampsia. Hypertension 65: 1307–1315

Janus P, Kuś P, Jaksik R, Vydra N, Toma-Jonik A, Gramatyka M, Kurpas M, Kimmel M & Widłak W (2024) Transcriptional responses to direct and indirect TGFB1 stimulation in cancerous and noncancerous mammary epithelial cells. Cell Commun Signal 22: 522

Jeggle P, Callies C, Tarjus A, Fassot C, Fels J, Oberleithner H, Jaisser F & Kusche-Vihrog K (2013) Epithelial sodium channel stiffens the vascular endothelium in vitro and in Liddle mice. Hypertension 61: 1053–1059

Jehanno C, Le Goff P, Habauzit D, Le Page Y, Lecomte S, Lecluze E, Percevault F, Avner S, Métivier R, Michel D, et al (2022) Hypoxia and ERα transcriptional crosstalk is associated with endocrine resistance in breast cancer. Cancers (Basel) 14: 4934

Joosse SA, Hannemann J, Spötter J, Bauche A, Andreas A, Müller V & Pantel K (2012) Changes in keratin expression during metastatic progression of breast cancer: impact on the detection of circulating tumor cells. Clin Cancer Res 18: 993–1003

Joseph R, Orlov YL, Huss M, Sun W, Kong SL, Ukil L, Pan YF, Li G, Lim M, Thomsen JS, et al (2010) Integrative model of genomic factors for determining binding site selection by estrogen receptor-α. Mol Syst Biol 6: 456

Kealy D, Ellerington R, Bansal S, Zeng AGX, Medeiros JJF, West KA, Blacknell N-M, Hawley CA, Lukaszonek J, Gawne RT, et al (2024) HIF-1 activated by PIM1 assembles a pathological transcription complex and regulon that drives JAK2V617F MPN disease. Cancer Biology

Koh MY & Powis G (2012) Passing the baton: the HIF switch. Trends Biochem Sci 37: 364–372

Ko J-H, Ko EA, Gu W, Lim I, Bang H & Zhou T (2013) Expression profiling of ion channel genes predicts clinical outcome in breast cancer. Mol Cancer 12: 106

Kremer D, Boddy K, Brown JJ, Davies DL, Fraser R, Lever AF, Morton JJ & Robertson JI (1977) Amiloride in the treatment of primary hyperaldosteronism and essential hypertension. Clin Endocrinol (Oxf) 7: 151–157

Laderoute KR (2005) The interaction between HIF-1 and AP-1 transcription factors in response to low oxygen. Semin Cell Dev Biol 16: 502–513

Lánczky A & Győrffy B (2021) Web-Based Survival Analysis Tool Tailored for Medical Research (KMplot): Development and Implementation. J Med Internet Res 23: e27633

Lando D, Peet DJ, Whelan DA, Gorman JJ & Whitelaw ML (2002) Asparagine hydroxylation of the HIF transactivation domain a hypoxic switch. Science 295: 858–861

Lawson H, Holt-Martyn JP, Dembitz V, Kabayama Y, Wang LM, Bellani A, Atwal S, Saffoon N, Durko J, van de Lagemaat LN, et al (2024) The selective prolyl hydroxylase inhibitor IOX5 stabilizes HIF-1α and compromises development and progression of acute myeloid leukemia. Nat Cancer 5: 916–937

Leslie TK, James AD, Zaccagna F, Grist JT, Deen S, Kennerley A, Riemer F, Kaggie JD, Gallagher FA, Gilbert FJ, et al (2019) Sodium homeostasis in the tumour microenvironment. Biochim Biophys Acta Rev Cancer 1872: 188304

Li H, Sun X, Li J, Liu W, Pan G, Mao A, Liu J, Zhang Q, Rao L, Xie X, et al (2022) Hypoxia induces docetaxel resistance in triple-negative breast cancer via the HIF-1α/miR-494/Survivin signaling pathway. Neoplasia 32: 100821

Lindström LS, Yau C, Czene K, Thompson CK, Hoadley KA, Van’t Veer LJ, Balassanian R, Bishop JW, Carpenter PM, Chen Y-Y, et al (2018) Intratumor Heterogeneity of the Estrogen Receptor and the Long-term Risk of Fatal Breast Cancer. J Natl Cancer Inst 110: 726–733

Liu Q, Palmgren VAC, Danen EH & Le Dévédec SE (2022a) Acute vs. chronic vs. intermittent hypoxia in breast Cancer: a review on its application in in vitro research. Mol Biol Rep 49: 10961–10973

Liu Z, Lee H-L, Suh JS, Deng P, Lee C-R, Bezouglaia O, Mirnia M, Chen V, Zhou M, Cui Z-K, et al (2022b) The ERα/KDM6B regulatory axis modulates osteogenic differentiation in human mesenchymal stem cells. Bone Res 10: 3

Loenarz C & Schofield CJ (2011) Physiological and biochemical aspects of hydroxylations and demethylations catalyzed by human 2-oxoglutarate oxygenases. Trends Biochem Sci 36: 7–18

Lombardi O, Li R, Halim S, Choudhry H, Ratcliffe PJ & Mole DR (2022) Pan-cancer analysis of tissue and single-cell HIF-pathway activation using a conserved gene signature. Cell Rep 41: 111652

Love MI, Huber W & Anders S (2014) Moderated estimation of fold change and dispersion for RNA-seq data with DESeq2. bioRxiv

Madak-Erdogan Z & Katzenellenbogen BS (2012) Aryl hydrocarbon receptor modulation of estrogen receptor α-mediated gene regulation by a multimeric chromatin complex involving the two receptors and the coregulator RIP140. Toxicol Sci 125: 401–411

Malcolm JR, Sajjaboontawee N, Yerlikaya S, Plunkett-Jones C, Boxall PJ & Brackenbury WJ (2023) Voltage-gated sodium channels, sodium transport and progression of solid tumours. Curr Top Membr 92: 71–98

Masters JR, Thomson JA, Daly-Burns B, Reid YA, Dirks WG, Packer P, Toji LH, Ohno T, Tanabe H, Arlett CF, et al (2001) Short tandem repeat profiling provides an international reference standard for human cell lines. Proc Natl Acad Sci U S A 98: 8012–8017

Matthews J & Gustafsson J-A (2006) Estrogen receptor and aryl hydrocarbon receptor signaling pathways. Nucl Recept Signal 4: e016

Matthews J, Wihlén B, Thomsen J & Gustafsson J-A (2005) Aryl hydrocarbon receptor-mediated transcription: ligand-dependent recruitment of estrogen receptor alpha to 2,3,7,8-tetrachlorodibenzo-p-dioxin-responsive promoters. Mol Cell Biol 25: 5317–5328

McQueen SRA, Chin WQ, Cunliffe HE & McDonald FJ (2025) Stable overexpression of the epithelial sodium channel alpha subunit reduces migration and proliferation in breast cancer cells. Breast Cancer Res Treat 211: 595–604

Melvin A & Rocha S (2012) Chromatin as an oxygen sensor and active player in the hypoxia response. Cell Signal 24: 35–43

Meyer AS, Miller MA, Gertler FB & Lauffenburger DA (2013) The receptor AXL diversifies EGFR signaling and limits the response to EGFR-targeted inhibitors in triple-negative breast cancer cells. Sci Signal 6: ra66

Miar A, Arnaiz E, Bridges E, Beedie S, Cribbs AP, Downes DJ, Beagrie RA, Rehwinkel J & Harris AL (2020) Hypoxia induces transcriptional and translational downregulation of the type I IFN pathway in multiple cancer cell types. Cancer Res 80: 5245–5256

Mohammed H, D’Santos C, Serandour AA, Ali HR, Brown GD, Atkins A, Rueda OM, Holmes KA, Theodorou V, Robinson JLL, et al (2013) Endogenous purification reveals GREB1 as a key estrogen receptor regulatory factor. Cell Rep 3: 342–349

Moriai R, Tsuji N, Moriai M, Kobayashi D & Watanabe N (2009) Survivin plays as a resistant factor against tamoxifen-induced apoptosis in human breast cancer cells. Breast Cancer Res Treat 117: 261–271

Morotti M, Bridges E, Valli A, Choudhry H, Sheldon H, Wigfield S, Gray N, Zois CE, Grimm F, Jones D, et al (2019) Hypoxia-induced switch in SNAT2/SLC38A2 regulation generates endocrine resistance in breast cancer. Proc Natl Acad Sci U S A 116: 12452–12461

Muz B, de la Puente P, Azab F & Azab AK (2015) The role of hypoxia in cancer progression, angiogenesis, metastasis, and resistance to therapy. Hypoxia (Auckl) 3: 83–92

Mylonis I, Chachami G, Samiotaki M, Panayotou G, Paraskeva E, Kalousi A, Georgatsou E, Bonanou S & Simos G (2006) Identification of MAPK phosphorylation sites and their role in the localization and activity of hypoxia-inducible factor-1alpha. J Biol Chem 281: 33095–33106

Nalwoga H, Ahmed L, Arnes JB, Wabinga H & Akslen LA (2016) Strong expression of hypoxia-inducible factor-1α (HIF-1α) is associated with Axl expression and features of aggressive tumors in African breast cancer. PLoS One 11: e0146823

Noreng S, Posert R, Bharadwaj A, Houser A & Baconguis I (2020) Molecular principles of assembly, activation, and inhibition in epithelial sodium channel. Elife 9: e59038

Ortmann BM (2024) Hypoxia-inducible factor in cancer: from pathway regulation to therapeutic opportunity. BMJ Oncol 3: e000154

Oxlund CS, Buhl KB, Jacobsen IA, Hansen MR, Gram J, Henriksen JE, Schousboe K, Tarnow L & Jensen BL (2014) Amiloride lowers blood pressure and attenuates urine plasminogen activation in patients with treatment-resistant hypertension. J Am Soc Hypertens 8: 872–881

Padró M, Louie RJ, Lananna BV, Krieg AJ, Timmerman LA & Chan DA (2017) Genome-independent hypoxic repression of estrogen receptor alpha in breast cancer cells. BMC Cancer 17: 203

Pan H, Gray R, Braybrooke J, Davies C, Taylor C, McGale P, Peto R, Pritchard KI, Bergh J, Dowsett M, et al (2017) 20-Year Risks of Breast-Cancer Recurrence after Stopping Endocrine Therapy at 5 Years. N Engl J Med 377: 1836–1846

Perillo B, Di Santi A, Cernera G, Ombra MN, Castoria G & Migliaccio A (2014) Nuclear receptor-induced transcription is driven by spatially and timely restricted waves of ROS. The role of Akt, IKKα, and DNA damage repair enzymes. Nucleus 5: 482–491

Perillo B, Tramontano A, Pezone A & Migliaccio A (2020) LSD1: more than demethylation of histone lysine residues. Exp Mol Med 52: 1936–1947

Place AE, Jin Huh S & Polyak K (2011) The microenvironment in breast cancer progression: biology and implications for treatment. Breast Cancer Res 13: 227

Premkumar DR, Adhikary G, Overholt JL, Simonson MS, Cherniack NS & Prabhakar NR (2000) Intracellular pathways linking hypoxia to activation of c-fos and AP-1. Adv Exp Med Biol 475: 101–109

Prickaerts P, Adriaens ME, van den Beucken T, Koch E, Dubois L, Dahlmans VEH, Gits C, Evelo CTA, Chan-Seng-Yue M, Wouters BG, et al (2016) Hypoxia increases genome-wide bivalent epigenetic marking by specific gain of H3K27me3. Epigenetics Chromatin 9: 46

Qadri YJ, Rooj AK & Fuller CM (2012) ENaCs and ASICs as therapeutic targets. Am J Physiol Cell Physiol 302: C943–65

Rae JM, Johnson MD, Scheys JO, Cordero KE, Larios JM & Lippman ME (2005) GREB 1 is a critical regulator of hormone dependent breast cancer growth. Breast Cancer Res Treat 92: 141–149

Ramírez F, Ryan DP, Grüning B, Bhardwaj V, Kilpert F, Richter AS, Heyne S, Dündar F & Manke T (2016) deepTools2: a next generation web server for deep-sequencing data analysis. Nucleic Acids Res 44: W160–5

Ribieras S, Tomasetto C & Rio MC (1998) The pS2/TFF1 trefoil factor, from basic research to clinical applications. Biochim Biophys Acta 1378: F61–77

Rogers ZJ, Colombani T, Khan S, Bhatt K, Nukovic A, Zhou G, Woolston BM, Taylor CT, Gilkes DM, Slavov N, et al (2024) Controlling pericellular oxygen tension in cell culture reveals distinct breast cancer responses to low oxygen tensions. Adv Sci (Weinh) 11: e2402557

Ross-Innes CS, Stark R, Holmes KA, Schmidt D, Spyrou C, Russell R, Massie CE, Vowler SL, Eldridge M & Carroll JS (2010) Cooperative interaction between retinoic acid receptor-alpha and estrogen receptor in breast cancer. Genes Dev 24: 171–182

Ross-Innes CS, Stark R, Teschendorff AE, Holmes KA, Ali HR, Dunning MJ, Brown GD, Gojis O, Ellis IO, Green AR, et al (2012) Differential oestrogen receptor binding is associated with clinical outcome in breast cancer. Nature 481: 389–393

Rozeboom B, Dey N & De P (2019) ER+ metastatic breast cancer: past, present, and a prescription for an apoptosis-targeted future. Am J Cancer Res 9: 2821–2831

Schödel J, Oikonomopoulos S, Ragoussis J, Pugh CW, Ratcliffe PJ & Mole DR (2011) High-resolution genome-wide mapping of HIF-binding sites by ChIP-seq. Blood 117: e207–17

Schofield CJ & Ratcliffe PJ (2004) Oxygen sensing by HIF hydroxylases. Nat Rev Mol Cell Biol 5: 343–354

Seward DJ, Cubberley G, Kim S, Schonewald M, Zhang L, Tripet B & Bentley DL (2007) Demethylation of trimethylated histone H3 Lys4 in vivo by JARID1 JmjC proteins. Nat Struct Mol Biol 14: 240–242

Shakir D, Batie M, Kwok C-S, Cook SJ, Kenneth NS & Rocha S (2025) NF-κB is a Central Regulator of Hypoxia-Induced Gene Expression. Molecular Biology

Shimkets RA, Warnock DG, Bositis CM, Nelson-Williams C, Hansson JH, Schambelan M, Gill JR Jr, Ulick S, Milora RV & Findling JW (1994) Liddle’s syndrome: heritable human hypertension caused by mutations in the beta subunit of the epithelial sodium channel. Cell 79: 407–414

Shin S & Janknecht R (2007) Diversity within the JMJD2 histone demethylase family. Biochem Biophys Res Commun 353: 973–977

Shi Y, Lan F, Matson C, Mulligan P, Whetstine JR, Cole PA, Casero RA & Shi Y (2004) Histone demethylation mediated by the nuclear amine oxidase homolog LSD1. Cell 119: 941–953

Siegl D, Kruchem M, Jansky S, Eichler E, Thies D, Hartwig U, Schuppan D & Bockamp E (2023) A PCR protocol to establish standards for routine mycoplasma testing that by design detects over ninety percent of all known mycoplasma species. iScience 26: 106724

Song Y, Dagil L, Fairall L, Robertson N, Wu M, Ragan TJ, Savva CG, Saleh A, Morone N, Kunze MBA, et al (2020) Mechanism of crosstalk between the LSD1 demethylase and HDAC1 deacetylase in the CoREST complex. Cell Rep 30: 2699–2711.e8

Spadazzi C, Mercatali L, Esposito M, Wei Y, Liverani C, De Vita A, Miserocchi G, Carretta E, Zanoni M, Cocchi C, et al (2021) Trefoil factor-1 upregulation in estrogen-receptor positive breast cancer correlates with an increased risk of bone metastasis. Bone 144: 115775

Strautnieks SS, Thompson RJ, Gardiner RM & Chung E (1996) A novel splice-site mutation in the gamma subunit of the epithelial sodium channel gene in three pseudohypoaldosteronism type 1 families. Nat Genet 13: 248–250

Tanimoto K, Makino Y, Pereira T & Poellinger L (2000) Mechanism of regulation of the hypoxia-inducible factor-1 alpha by the von Hippel-Lindau tumor suppressor protein. EMBO J 19: 4298–4309

Teiwes J & Toto RD (2007) Epithelial sodium channel inhibition in cardiovascular disease. A potential role for amiloride. Am J Hypertens 20: 109–117

Tsukada Y-I, Fang J, Erdjument-Bromage H, Warren ME, Borchers CH, Tempst P & Zhang Y (2006) Histone demethylation by a family of JmjC domain-containing proteins. Nature 439: 811–816

Turner KJ, Crew JP, Wykoff CC, Watson PH, Poulsom R, Pastorek J, Ratcliffe PJ, Cranston D & Harris AL (2002) The hypoxia-inducible genes VEGF and CA9 are differentially regulated in superficial vs invasive bladder cancer. Br J Cancer 86: 1276–1282

Tutzauer J, Sjöström M, Holmberg E, Karlsson P, Killander F, Leeb-Lundberg LMF, Malmström P, Niméus E, Fernö M & Jögi A (2022) Breast cancer hypoxia in relation to prognosis and benefit from radiotherapy after breast-conserving surgery in a large, randomised trial with long-term follow-up. Br J Cancer 126: 1145–1156

Uphoff CC, Gignac SM & Drexler HG (1992) Mycoplasma contamination in human leukemia cell lines. I. Comparison of various detection methods. J Immunol Methods 149: 43–53

Wang J, Wang Y, Duan Z & Hu W (2020) Hypoxia-induced alterations of transcriptome and chromatin accessibility in HL-1 cells. IUBMB Life 72: 1737–1746

Wang L, Xu C, Liu X, Yang Y, Cao L, Xiang G, Liu F, Wang S, Liu J, Meng Q, et al (2019) TGF-β1 stimulates epithelial-mesenchymal transition and cancer-associated myoepithelial cell during the progression from in situ to invasive breast cancer. Cancer Cell Int 19: 343

Wang P, Zhang X-P, Liu F & Wang W (2025) Progressive deactivation of hydroxylases controls hypoxia-inducible factor-1α-coordinated cellular adaptation to graded hypoxia. Research (Wash DC) 8: 0651

Ware AW, Harris JJ, Slatter TL, Cunliffe HE & McDonald FJ (2021) The epithelial sodium channel has a role in breast cancer cell proliferation. Breast Cancer Res Treat 187: 31–43

Wickham H (2016) Ggplot2 2nd ed. Basel, Switzerland: Springer International Publishing

Withnell E & Secrier M (2024) SpottedPy quantifies relationships between spatial transcriptomic hotspots and uncovers environmental cues of epithelial-mesenchymal plasticity in breast cancer. Genome Biol 25: 289

Wolff M, Kosyna FK, Dunst J, Jelkmann W & Depping R (2017) Impact of hypoxia inducible factors on estrogen receptor expression in breast cancer cells. Arch Biochem Biophys 613: 23–30

Wykoff CC, Beasley NJ, Watson PH, Turner KJ, Pastorek J, Sibtain A, Wilson GD, Turley H, Talks KL, Maxwell PH, et al (2000) Hypoxia-inducible expression of tumor-associated carbonic anhydrases. Cancer Res 60: 7075–7083

Xin J, Zhang H, He Y, Duren Z, Bai C, Chen L, Luo X, Yan D-S, Zhang C, Zhu X, et al (2020) Chromatin accessibility landscape and regulatory network of high-altitude hypoxia adaptation. Nat Commun 11: 4928

Xiong Z-G, Pignataro G, Li M, Chang S-Y & Simon RP (2008) Acid-sensing ion channels (ASICs) as pharmacological targets for neurodegenerative diseases. Curr Opin Pharmacol 8: 25–32

Xu W, Mezencev R, Kim B, Wang L, McDonald J & Sulchek T (2012) Cell stiffness is a biomarker of the metastatic potential of ovarian cancer cells. PLoS One 7: e46609

Yang H-Y, Charles R-P, Hummler E, Baines DL & Isseroff RR (2013) The epithelial sodium channel mediates the directionality of galvanotaxis in human keratinocytes. J Cell Sci 126: 1942–1951

Yang J, AlTahan A, Jones DT, Buffa FM, Bridges E, Interiano RB, Qu C, Vogt N, Li J-L, Baban D, et al (2015) Estrogen receptor-α directly regulates the hypoxia-inducible factor 1 pathway associated with antiestrogen response in breast cancer. Proc Natl Acad Sci U S A 112: 15172–15177

Yang J, Jubb AM, Pike L, Buffa FM, Turley H, Baban D, Leek R, Gatter KC, Ragoussis J & Harris AL (2010) The histone demethylase JMJD2B is regulated by estrogen receptor alpha and hypoxia, and is a key mediator of estrogen induced growth. Cancer Res 70: 6456–6466

Yu NY, Iftimi A, Yau C, Tobin NP, van’t Veer L, Hoadley KA, Benz CC, Nordenskjöld B, Fornander T, Stål O, et al (2019) Assessment of long-term distant recurrence-free survival associated with tamoxifen therapy in postmenopausal patients with luminal A or luminal B breast cancer. JAMA Oncol 5: 1304–1309

Zelzer E, Levy Y, Kahana C, Shilo BZ, Rubinstein M & Cohen B (1998) Insulin induces transcription of target genes through the hypoxia-inducible factor HIF-1alpha/ARNT. EMBO J 17: 5085–5094

Zhang Y, Liu T, Meyer CA, Eeckhoute J, Johnson DS, Bernstein BE, Nusbaum C, Myers RM, Brown M, Li W, et al (2008) Model-based analysis of ChIP-Seq (MACS). Genome Biol 9: R137

Zhao W, Rose SF, Blake R, Godicelj A, Cullen AE, Stenning J, Beevors L, Gehrung M, Kumar S, Kishore K, et al (2024) ZMIZ1 enhances ERα-dependent expression of E2F2 in breast cancer. J Mol Endocrinol 73

