## Supplementary Figures and Tables for "An ERα-Dependent Hypoxia Response Defines EMT-Adjacent Tumour Regions and Suppresses the Pro-survival Effects of Amiloride in Estrogen Receptor-Positive Breast Cancer"

#### Figure S1

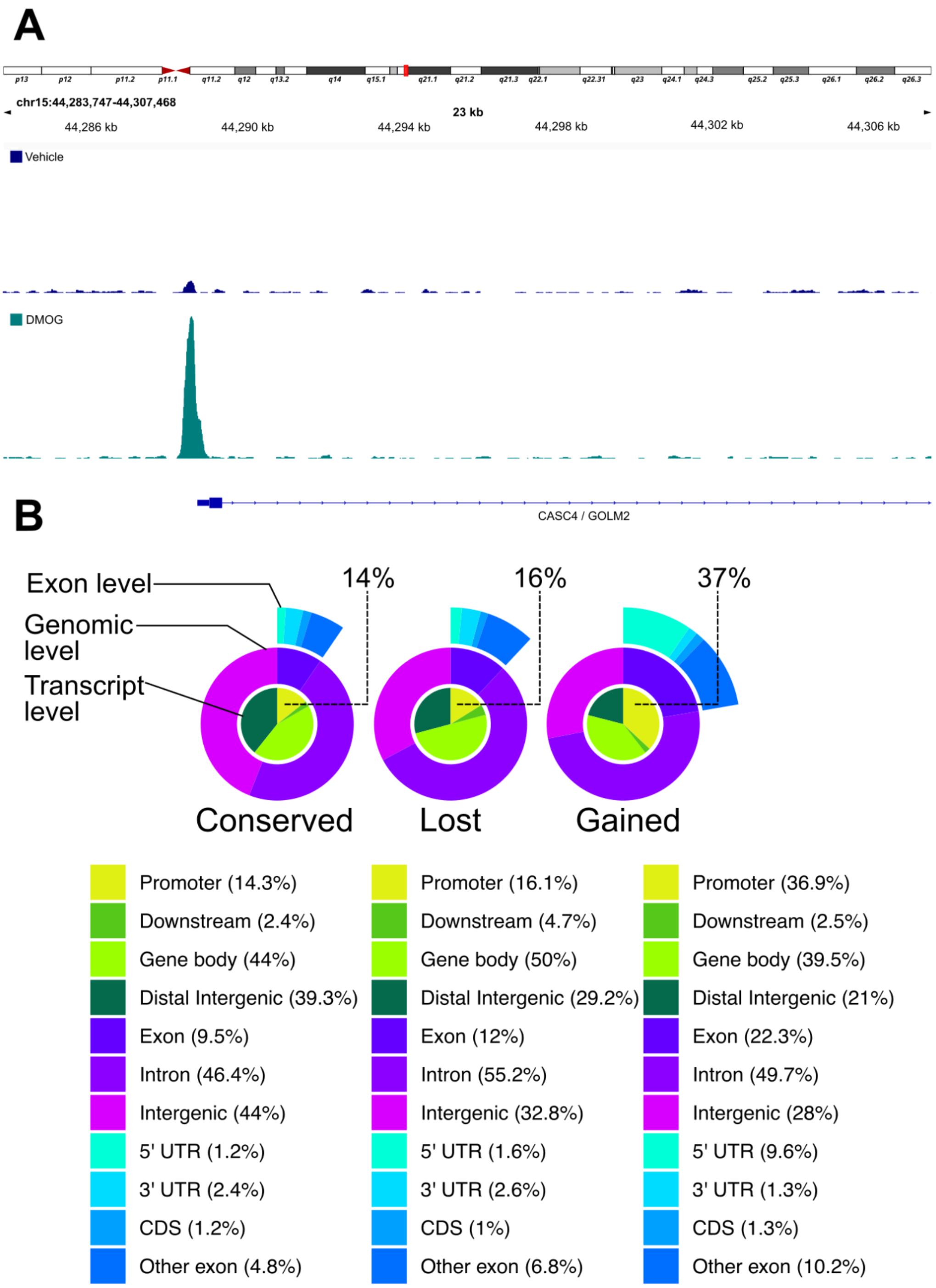


**Figure S1** (A) Individual analysis of *CASC4/GOLM2* promoter. (B) Expanded analysis of conserved, gained, and lost ERα ChIP-seq loci shows that the gained sites are enriched for promoter regions (37%) when compared to the other conserved and lost ERα binding sites (CDS = Coding Sequence).

### Figure S2

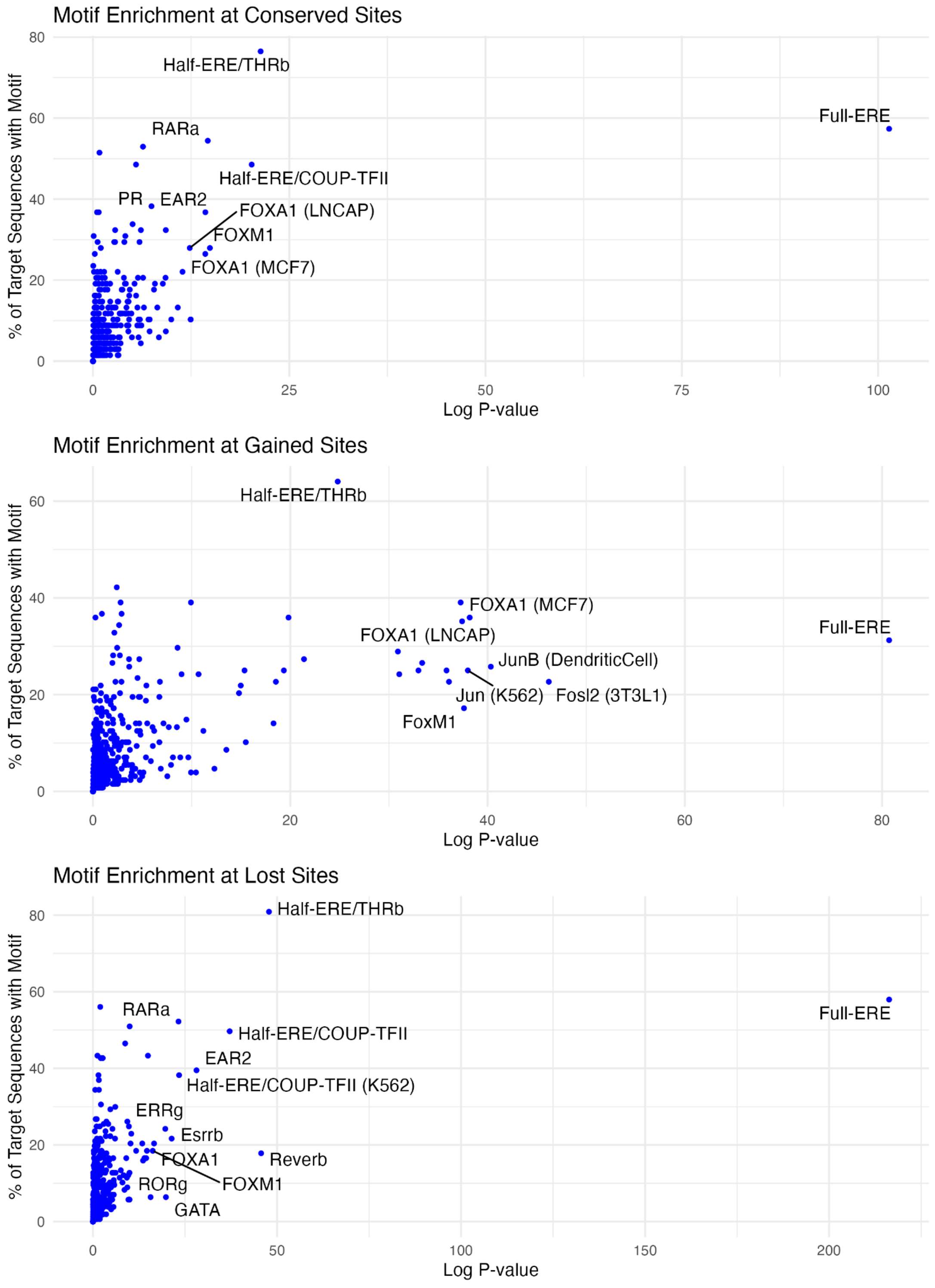


**Figure S2 Motif enrichment analysis of ERα binding sites following DMOG treatment.** Expanded motif analysis of ERα ChIP-seq peaks identified as conserved, gained, or lost after 16 h of DMOG exposure. Each panel shows the percentage of target sequences containing the indicated motif (y-axis) versus statistical significance (log-transformed p-value, x-axis). (Top) Conserved ERα binding sites are strongly enriched for classical estrogen response elements (EREs) and known ERα co-factors including FOXA1, RARα and COUP-TFII. (Middle) Gained sites retain ERE enrichment but show increased association with motifs for FOS (22%) and JunB (25%), as well as FOXA1. (Bottom) Lost sites also contain EREs and enrichment for motifs associated with nuclear receptors (e.g., RARα, EAR2(NR2F6)) along with FOXA1.

### Figure S3

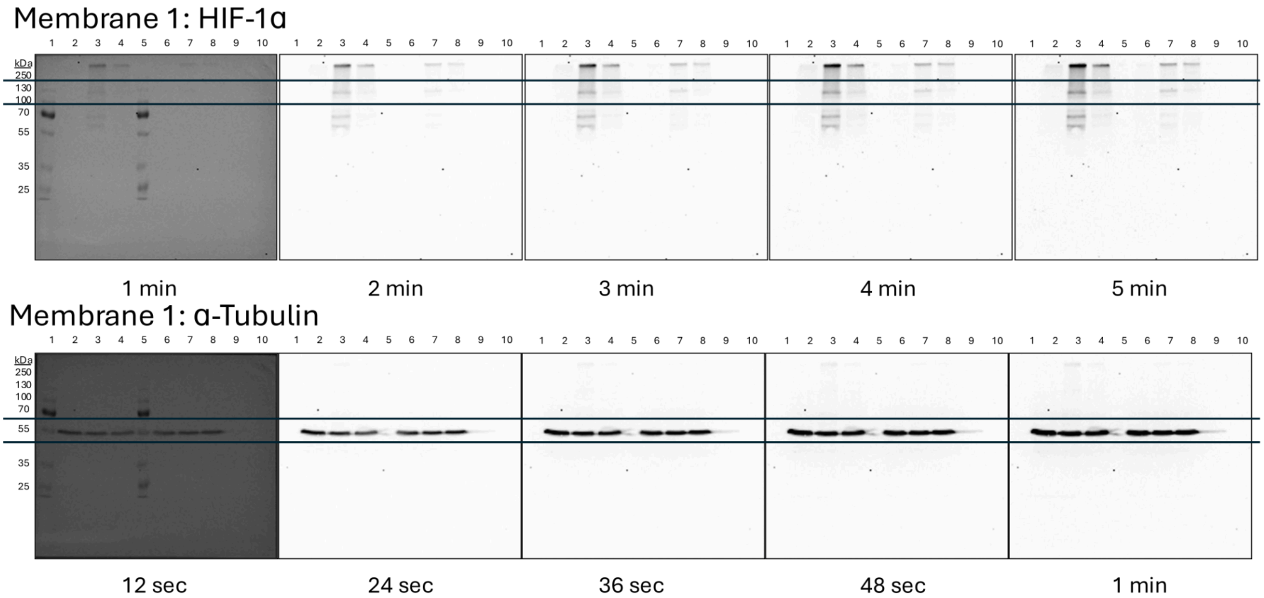


1. Ladder
2. MCF-7 0 hour (normoxia)
3. MCF-7 8 hour 1% $O_2$
4. MCF-7 48 hout 1% $O_2$
5. Ladder
6. T-47D 0 hour (normoxxia)
7. T-47D 8 hour 1% $O_2$
8. T-47D 48 hour 1% $O_2$
9. Blank
10. Blank

HIF-1α ~96 kDa
(CST 36169, 1:1,000)
2° 1:4000

α-Tubulin ~50 kDa
(Merck T6199, 1:5,000)
2° 1:5,000

**Figure S3 Full western blots corresponding to Figure 3A.** Uncropped western blot images for MCF7 and T47D cell lines probed for HIF-1α and under normoxia, and at 8 h and 48 h following hypoxic culture (1% $O_2$). These blots demonstrate induction of HIF-1α at 8 h with reduced signal at 48 h.

### Figure S4

**A**

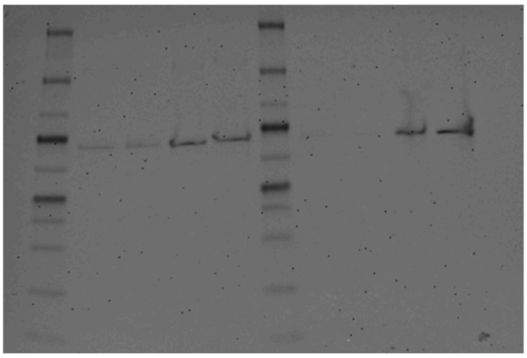

**B**

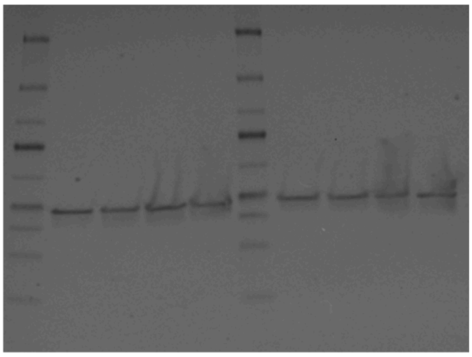

**Antibody Dilution**
**ER- alpha (Ab32063) 1:1000**
**GAPDH (10494-1-AP) 1:7500**
**Secondary anti-rabbit 1:10,000**

**A)** Western blot image of cell lysates probed with ER-alpha antibody. Lane 1 and lane 6 = Pre-stained marker (associated molecular weights shown 260kDa, 140kDa, 100kDa, 70kDa, 50kDa, 40kDa, 35kDa, 25kDa, 15KDa and 10kDa), Lane 2 and 3 = MCF7 cells treated with 100nM fulvestrant for 48 hours, Lane 4 and 5 = MCF7 cells treated with 0nM of fulvestrant (vehicle control) for 48 hours, lane 7 and 8 = MCF7 cells treated with 100nM fulvestrant for 96 hours, lane 9 and 10 = MCF7 treated with 0nM of fulvestrant (vehicle control) for 96 hours. NB predicted band size or ER alpha = 66kDa. Exposed for 10 minutes.

**B)** Western blot image of cell lysates probed with GAPDH antibody. Lane 1 and lane 6 = Pre-stained marker (associated molecular weights shown 260kDa, 140kDa, 100kDa, 70kDa, 50kDa, 40kDa, 35kDa, 25kDa, 15KDa and 10kDa), Lane 2 and 3 = MCF7 cells treated with 100nM fulvestrant for 46 hours, Lane 4 and 5 = MCF7 cells treated with 0nM of fulvestrant (vehicle control) for 48 hours, lane 7 and 8 = MCF7 cells treated with 100nM fulvestrant for 96 hours, lane 9 and 10 = MCF7 treated with 0nM of fulvestrant (vehicle control) for 96 hours. NB predicted band size or GAPDH = 36kDa. Exposed for 6 seconds

**Figure S4. Fulvestrant effectively depletes ERα protein in MCF7 cells under hypoxia. (A)** Western blot showing ERα protein levels in MCF7 cells treated with 100 nM fulvestrant or vehicle for 48 or 96 hours. **(B)** Corresponding GAPDH blot as a loading control (predicted size: 36 kDa). Antibodies: ERα (Ab32063); GAPDH (10494-1-AP).

Figure S5

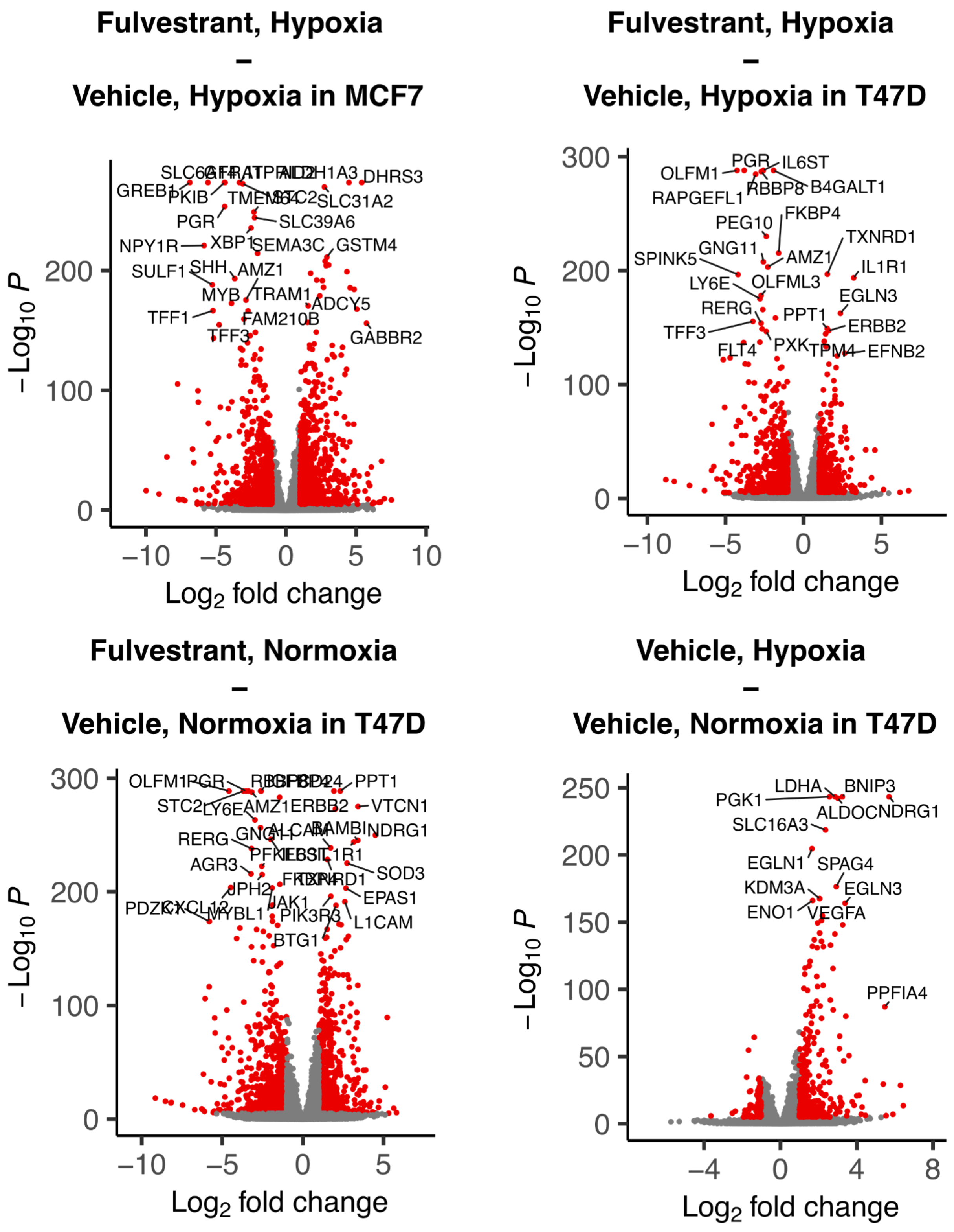
Fulvestrant, Hypoxia
–
Vehicle, Hypoxia in MCF7
Fulvestrant, Hypoxia
–
Vehicle, Hypoxia in T47D
Fulvestrant, Normoxia
–
Vehicle, Normoxia in T47D
Vehicle, Hypoxia
–
Vehicle, Normoxia in T47D
$-Log_{10}$ P
$Log_2$ fold change
GREB1
PKIB
PGR
NPY1R
XBP1
SULF1
SHH
AMZ1
MYB
TRAM1
TFF1
FAM210B
TFF3
DHRS3
SLC31A2
SLC39A6
SEMA3C
GSTM4
ADCY5
GABBR2
OLFM1
PGR
IL6ST
RAPGEFL1
RBBP8
B4GALT1
PEG10
FKBP4
TXNRD1
GNG11
SPINK5
AMZ1
IL1R1
LY6E
OLFML3
EGLN3
RERG
PPT1
ERBB2
TFF3
PXK
EFNB2
FLT4
PPT1
VTCN1
NDRG1
RERG
AGR3
SOD3
JPH2
EPAS1
JAK1
L1CAM
BTG1
LDHA
BNIP3
PGK1
SLC16A3
EGLN1
SPAG4
KDM3A
EGLN3
ENO1
VEGFA
PPFIA4

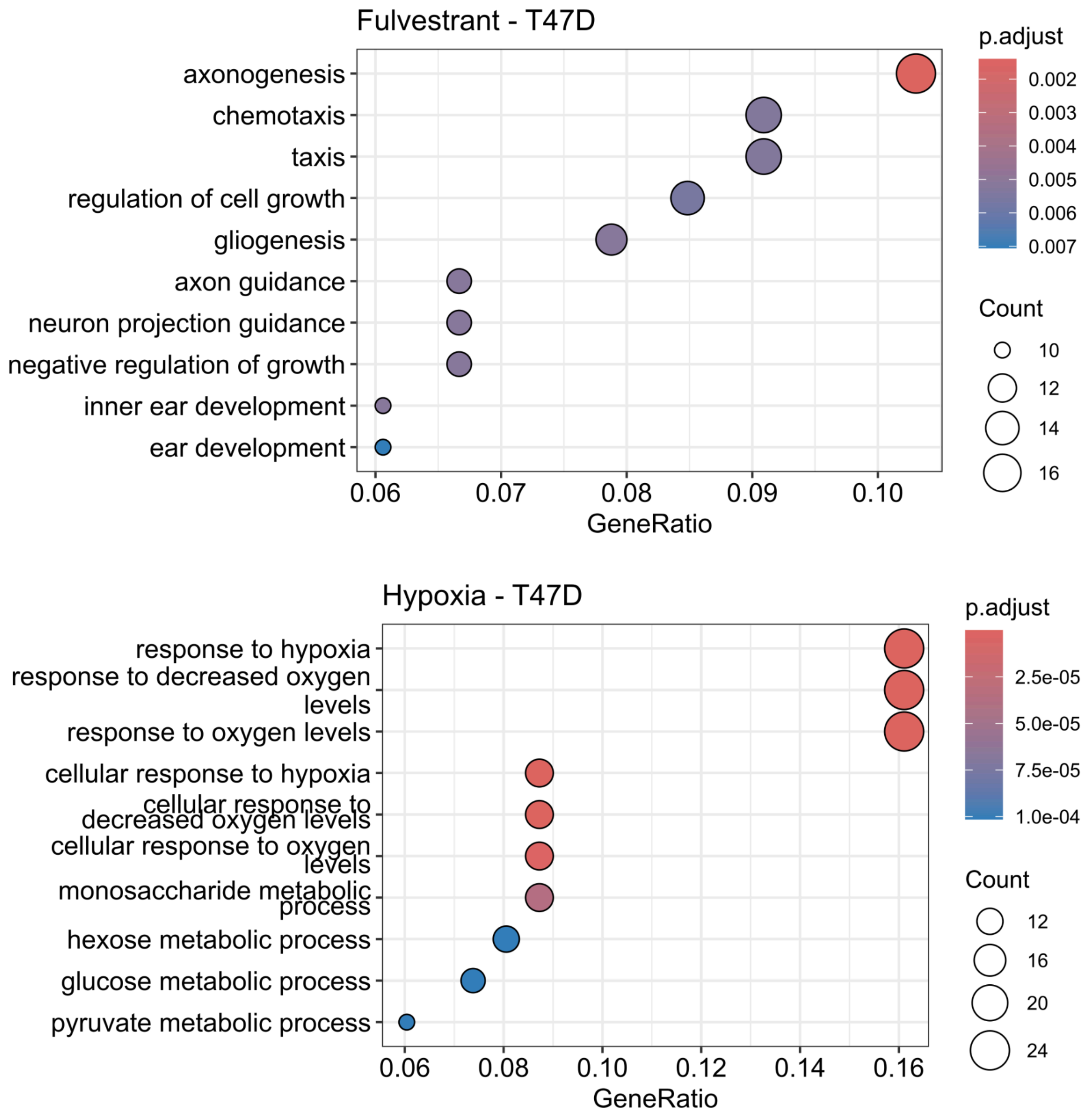


**Figure S5. Volcano plots and GO analysis of T47D transcriptomic responses.** Differential gene expression (DGE) and gene ontology (GO) enrichment in cells across different conditions. (Top) Volcano plots show DEGs for hypoxia ± fulvestrant in MCF7 cells. (Middle) Volcano plots show DEGs for fulvestrant and hypoxia and hypoxia alone in MCF7 cells. (Bottom) GO enrichment of DEGs following fulvestrant treatment shows suppression of growth and chemotaxis, and GO enrichment of DEGs following hypoxia treatment reveals upregulation of classical hypoxia-responsive and metabolic pathways.

Figure S6

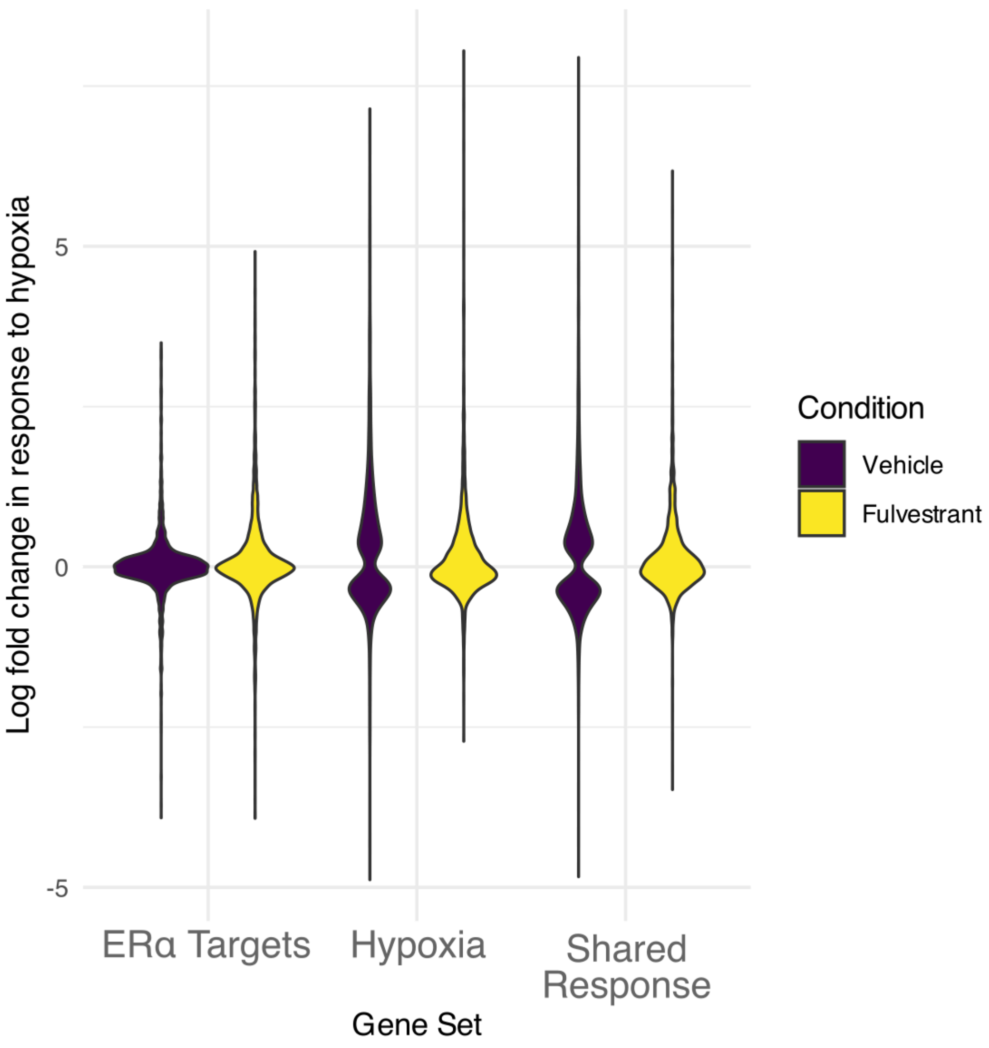


**Figure S6 Target ERα signalling dampens the signed LFC of the hypoxia response in MCF7 cells.** Analysis of our three transcriptomic gene sets demonstrated that ERα-specific targets were poorly responsive to hypoxia. In contrast, transcripts from our hypoxic gene set displayed a shift in the LFC in response to hypoxia towards zero following fulvestrant treatment. Similarly, our shared response gene set exhibited a comparable shift in the distribution of LFCs, along with a reduction in the maximum LFC values.

#### Figure S7

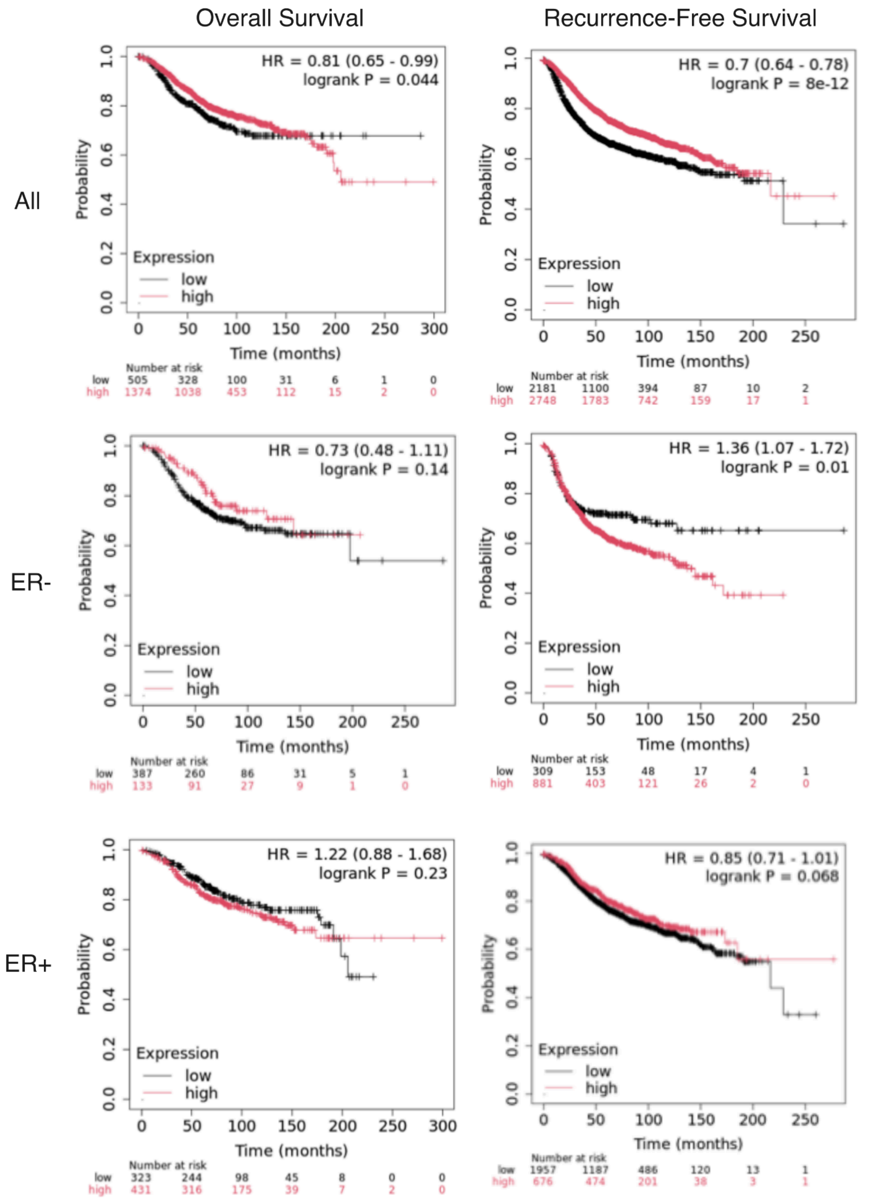


**Figure S7 Survival analysis based on SCNN1A expression.** Kaplan-Meier plots generated from Kmplot.com, showing overall survival and recurrence-free survival in breast cancer patients stratified by SCNN1A expression. Analyses are shown for the full cohort (All patients) and further stratified by ERα status.

### Table S1 Conserved ERα binding sites

| Chromosome | Start | End | Feature | Location | Distance (bp) |
|---|---|---|---|---|---|
| chr1 | 7447895 | 7448091 | ENSG00000225126 | upstream | 6341 |
| chr2 | 11539566 | 11539842 | ENSG00000264010 | downstream | 763 |
| chr2 | 11532400 | 11532646 | ENSG00000196208 | upstream | 1470 |
| chr2 | 237559046 | 237559273 | ENSG00000222449 | upstream | 3892 |
| chr2 | 11122452 | 11122608 | ENSG00000145063 | inside | 10213 |
| chr2 | 11498588 | 11498888 | ENSG00000201610 | upstream | 18509 |
| chr2 | 57140497 | 57140673 | ENSG00000276201 | downstream | 52063 |
| chr2 | 45752663 | 45752844 | ENSG00000228481 | upstream | 77187 |
| chr2 | 35297511 | 35297710 | ENSG00000196085 | upstream | 77907 |
| chr2 | 225843116 | 225843340 | ENSG00000279015 | downstream | 81004 |
| chr3 | 156812587 | 156812788 | ENSG00000230457 | upstream | 1855 |
| chr3 | 161339676 | 161339951 | ENSG00000243838 | upstream | 9438 |
| chr4 | 74661771 | 74661963 | ENSG00000249942 | upstream | 72290 |
| chr5 | 19735104 | 19735291 | ENSG00000264928 | downstream | 219 |
| chr5 | 118009980 | 118010173 | ENSG00000249797 | inside | 84972 |
| chr5 | 106241582 | 106241861 | ENSG00000251204 | downstream | 173715 |
| chr6 | 17393838 | 17394068 | ENSG00000112186 | inside | 622 |
| chr6 | 17389443 | 17389567 | ENSG00000112186 | upstream | 3649 |
| chr6 | 135211315 | 135211515 | ENSG00000236703 | upstream | 15320 |
| chr6 | 15221868 | 15222196 | ENSG00000271888 | downstream | 21727 |
| chr7 | 84536226 | 84536406 | ENSG00000235139 | inside | 3750 |
| chr7 | 45543559 | 45543770 | ENSG00000226999 | upstream | 8878 |
| chr7 | 867163 | 867528 | ENSG00000239857 | upstream | 9024 |
| chr8 | 68853422 | 68853605 | ENSG00000254337 | upstream | 659 |
| chr8 | 68854486 | 68854746 | ENSG00000254337 | upstream | 1723 |

| | | | | | |
|---|---|---|---|---|---|
| chr8 | 101558416 | 101558812 | ENSG00000253153 | downstream | 3196 |
| chr8 | 128141948 | 128142235 | ENSG00000221261 | upstream | 7881 |
| chr8 | 101543238 | 101543438 | ENSG00000253991 | upstream | 13669 |
| chr8 | 127870655 | 127870876 | ENSG00000278324 | upstream | 19750 |
| chr8 | 102600977 | 102601177 | ENSG00000253382 | downstream | 20332 |
| chr8 | 90887703 | 90887896 | ENSG00000254251 | upstream | 28463 |
| chr8 | 121151018 | 121151201 | ENSG00000221644 | downstream | 35557 |
| chr8 | 127668704 | 127668936 | ENSG00000279524 | upstream | 38047 |
| chr8 | 115867563 | 115867756 | ENSG00000104447 | upstream | 57890 |
| chr9 | 88530074 | 88530293 | ENSG00000130045 | upstream | 4808 |
| chr9 | 92618750 | 92618925 | ENSG00000127080 | inside | 5566 |
| chr9 | 112012183 | 112012386 | ENSG00000222356 | upstream | 15444 |
| chr9 | 108478875 | 108478958 | ENSG00000222512 | upstream | 119946 |
| chr10 | 8726078 | 8726301 | ENSG00000270234 | downstream | 1790 |
| chr10 | 69439412 | 69439615 | ENSG00000232734 | upstream | 6041 |
| chr10 | 8128395 | 8128615 | ENSG00000225053 | upstream | 32645 |
| chr10 | 93435121 | 93435441 | ENSG00000138119 | inside | 46876 |
| chr10 | 115664751 | 115665018 | ENSG00000270965 | downstream | 154824 |
| chr10 | 29124951 | 29125259 | ENSG00000120563 | upstream | 163802 |
| chr11 | 118436223 | 118436443 | ENSG00000118058 | upstream | 47 |
| chr11 | 65418756 | 65418952 | ENSG00000245532 | upstream | 3822 |
| chr11 | 86776215 | 86776447 | ENSG00000150687 | upstream | 14612 |
| chr11 | 79059982 | 79060212 | ENSG00000255345 | upstream | 32636 |
| chr11 | 30521748 | 30521977 | ENSG00000254489 | upstream | 62153 |
| chr11 | 101336289 | 101336469 | ENSG00000263885 | downstream | 183351 |
| chr12 | 2795961 | 2796178 | ENSG00000004478 | inside | 1008 |
| chr12 | 116390487 | 116390710 | ENSG00000258346 | downstream | 1016 |
| chr12 | 97568787 | 97568990 | ENSG00000277081 | downstream | 4460 |

| | | | | | |
|---|---|---|---|---|---|
| chr12 | 75706476 | 75706664 | ENSG00000258088 | upstream | 7660 |
| chr12 | 52895270 | 52895464 | ENSG00000241146 | downstream | 8885 |
| chr12 | 121748867 | 121749168 | ENSG00000188735 | inside | 33833 |
| chr12 | 89465042 | 89465259 | ENSG00000226982 | upstream | 34834 |
| chr12 | 74871857 | 74872153 | ENSG00000257998 | upstream | 143006 |
| chr13 | 68686120 | 68686322 | ENSG00000243671 | downstream | 7482 |
| chr14 | 91276509 | 91276715 | ENSG00000260810 | downstream | 17506 |
| chr14 | 67480113 | 67480315 | ENSG00000265993 | upstream | 38183 |
| chr14 | 33781446 | 33781659 | ENSG00000265244 | downstream | 195651 |
| chr15 | 44288466 | 44288642 | ENSG00000166734 | upstream | 87 |
| chr15 | 92645621 | 92645767 | ENSG00000275965 | downstream | 10956 |
| chr15 | 98749253 | 98749463 | ENSG00000264480 | upstream | 34963 |
| chr15 | 95894432 | 95894661 | ENSG00000275443 | upstream | 95921 |
| chr16 | 14370277 | 14370502 | ENSG00000260009 | upstream | 11 |
| chr16 | 85306204 | 85306396 | ENSG00000266307 | includeFeature | 22 |
| chr16 | 85466018 | 85466265 | ENSG00000278716 | upstream | 23548 |
| chr17 | 63916679 | 63917305 | ENSG00000259384 | overlapEnd | 105 |
| chr17 | 40448715 | 40448982 | ENSG00000141753 | inside | 5254 |
| chr17 | 57899911 | 57900078 | ENSG00000264914 | upstream | 12255 |
| chr17 | 66952285 | 66952478 | ENSG00000075461 | upstream | 12432 |
| chr19 | 40921208 | 40921423 | ENSG00000256612 | upstream | 2796 |
| chr19 | 16463991 | 16464213 | ENSG00000127527 | inside | 7872 |
| chr20 | 60062974 | 60063333 | ENSG00000176659 | inside | 7049 |
| chr20 | 48864805 | 48865063 | ENSG00000278231 | upstream | 15347 |
| chr20 | 24649942 | 24650269 | ENSG00000231015 | upstream | 29278 |
| chr20 | 54123649 | 54123967 | ENSG00000019186 | downstream | 29482 |
| chr20 | 48572298 | 48572473 | ENSG00000238452 | upstream | 77989 |
| chr21 | 34734779 | 34734955 | ENSG00000230978 | inside | 2226 |

| | | | | | |
|---|---|---|---|---|---|
| chr21 | 40324568 | 40325001 | ENSG00000235123 | upstream | 58082 |
| chr21 | 15210333 | 15210598 | ENSG00000280082 | upstream | 136054 |
| chrX | 150041314 | 150041484 | ENSG00000183171 | upstream | 72894 |

### Table S2 Lost ERα binding sites

| Chromosome | Start | End | Feature | Location | Distance (bp) |
|---|---|---|---|---|---|
| chr1 | 1073738 | 1073916 | ENSG00000237330 | inside | 391 |
| chr1 | 1739505 | 1739703 | ENSG00000268575 | upstream | 1817 |
| chr1 | 15229919 | 15230116 | ENSG00000278480 | downstream | 2370 |
| chr1 | 42992481 | 42992665 | ENSG00000207256 | downstream | 937 |
| chr1 | 108237382 | 108237572 | ENSG00000196427 | inside | 6509 |
| chr1 | 108382300 | 108382541 | ENSG00000243967 | inside | 6462 |
| chr1 | 108456758 | 108456986 | ENSG00000238118 | upstream | 35463 |
| chr1 | 109240176 | 109240364 | ENSG00000143126 | upstream | 9655 |
| chr1 | 111513440 | 111513829 | ENSG00000200360 | upstream | 23017 |
| chr1 | 145708230 | 145708485 | ENSG00000174827 | upstream | 82 |
| chr1 | 162159502 | 162159739 | ENSG00000266144 | downstream | 2320 |
| chr1 | 200016853 | 200017168 | ENSG00000202329 | upstream | 2058 |
| chr1 | 202112905 | 202113111 | ENSG00000170075 | upstream | 9747 |
| chr1 | 203089359 | 203089683 | ENSG00000163485 | upstream | 971 |
| chr1 | 206903702 | 206903979 | ENSG00000271680 | upstream | 1949 |
| chr1 | 234963263 | 234963475 | ENSG00000237520 | downstream | 3274 |
| chr2 | 11531036 | 11531233 | ENSG00000196208 | upstream | 2883 |
| chr2 | 33070225 | 33070427 | ENSG00000223951 | upstream | 7112 |
| chr2 | 39180352 | 39180563 | ENSG00000205111 | inside | 4706 |
| chr2 | 43273644 | 43273847 | ENSG00000234936 | downstream | 40250 |
| chr2 | 99636693 | 99636968 | ENSG00000222274 | downstream | 116461 |
| chr2 | 138435798 | 138435979 | ENSG00000232915 | upstream | 22480 |
| chr2 | 158723550 | 158723750 | ENSG00000204380 | inside | 11252 |
| chr2 | 176845788 | 176845963 | ENSG00000227098 | downstream | 30 |

| | | | | | |
|---|---|---|---|---|---|
| chr2 | 185187415 | 185187662 | ENSG00000237824 | upstream | 284853 |
| chr2 | 217247531 | 217247724 | ENSG00000251982 | upstream | 3832 |
| chr2 | 217388117 | 217388332 | ENSG00000278004 | downstream | 104594 |
| chr2 | 236769857 | 236770035 | ENSG00000232328 | downstream | 15494 |
| chr2 | 237041374 | 237041592 | ENSG00000124835 | downstream | 7007 |
| chr2 | 237496478 | 237496697 | ENSG00000115648 | inside | 11050 |
| chr2 | 240069934 | 240070143 | ENSG00000231278 | upstream | 9290 |
| chr3 | 15292889 | 15293087 | ENSG00000224660 | downstream | 28396 |
| chr3 | 53145340 | 53145539 | ENSG00000163932 | upstream | 10470 |
| chr3 | 64491486 | 64491689 | ENSG00000212340 | upstream | 20066 |
| chr3 | 100104194 | 100104379 | ENSG00000168386 | inside | 10134 |
| chr3 | 107999532 | 107999870 | ENSG00000279277 | downstream | 36848 |
| chr3 | 134327693 | 134327869 | ENSG00000240006 | downstream | 6522 |
| chr3 | 150737280 | 150737622 | ENSG00000244668 | upstream | 16444 |
| chr3 | 161272679 | 161272894 | ENSG00000271052 | upstream | 49532 |
| chr3 | 189842710 | 189843004 | ENSG00000216058 | downstream | 12701 |
| chr3 | 193879613 | 193879823 | ENSG00000229155 | upstream | 35589 |
| chr3 | 194159026 | 194159283 | ENSG00000242201 | downstream | 13055 |
| chr3 | 194495083 | 194495263 | ENSG00000133657 | inside | 3101 |
| chr4 | 6415384 | 6415591 | ENSG00000109501 | downstream | 112119 |
| chr4 | 38462665 | 38462875 | ENSG00000249667 | upstream | 46892 |
| chr4 | 67928684 | 67928922 | ENSG00000187054 | inside | 19299 |
| chr4 | 102404910 | 102405092 | ENSG00000138821 | inside | 26166 |
| chr5 | 129177721 | 129177911 | ENSG00000264563 | downstream | 79955 |
| chr5 | 132378677 | 132378896 | ENSG00000233006 | upstream | 8761 |
| chr5 | 139643491 | 139643676 | ENSG00000272255 | downstream | 784 |
| chr5 | 168316437 | 168316664 | ENSG00000113645 | inside | 24786 |
| chr5 | 173472727 | 173473025 | ENSG00000253141 | inside | 9227 |

| | | | | | |
|---|---|---|---|---|---|
| chr5 | 174483505 | 174483684 | ENSG00000213376 | upstream | 29621 |
| chr6 | 1641363 | 1641589 | ENSG00000054598 | downstream | 27466 |
| chr6 | 15036802 | 15036980 | ENSG00000234261 | inside | 53023 |
| chr6 | 44110446 | 44110621 | ENSG00000137216 | upstream | 16293 |
| chr6 | 85272042 | 85272224 | ENSG00000217060 | upstream | 13852 |
| chr6 | 110555058 | 110555234 | ENSG00000219559 | downstream | 6941 |
| chr6 | 125202306 | 125202518 | ENSG00000111907 | inside | 61889 |
| chr6 | 125584729 | 125584990 | ENSG00000224506 | upstream | 89363 |
| chr6 | 134117327 | 134117587 | ENSG00000216753 | upstream | 1591 |
| chr6 | 138582615 | 138582795 | ENSG00000216802 | downstream | 67431 |
| chr6 | 149080270 | 149080470 | ENSG00000219487 | downstream | 31539 |
| chr7 | 2689188 | 2689397 | ENSG00000174945 | inside | 9666 |
| chr7 | 16880631 | 16880806 | ENSG00000173467 | inside | 1181 |
| chr7 | 80942981 | 80943166 | ENSG00000075223 | upstream | 20622 |
| chr7 | 113281408 | 113281604 | ENSG00000277061 | upstream | 63866 |
| chr7 | 117124598 | 117124837 | ENSG00000274344 | downstream | 6615 |
| chr7 | 140396582 | 140396772 | ENSG00000146955 | upstream | 7271 |
| chr7 | 151856632 | 151856876 | ENSG00000239911 | upstream | 20166 |
| chr8 | 17329743 | 17329939 | ENSG00000264599 | downstream | 25708 |
| chr8 | 22739166 | 22739363 | ENSG00000179388 | upstream | 45864 |
| chr8 | 37062296 | 37062498 | ENSG00000254038 | downstream | 4943 |
| chr8 | 67381088 | 67381268 | ENSG00000241961 | upstream | 25247 |
| chr8 | 85840889 | 85841129 | ENSG00000253154 | inside | 1184 |
| chr8 | 94092652 | 94092837 | ENSG00000253585 | downstream | 4927 |
| chr8 | 97764221 | 97764443 | ENSG00000202399 | upstream | 7870 |
| chr8 | 101466743 | 101466946 | ENSG00000254084 | upstream | 14291 |
| chr8 | 127800785 | 127801057 | ENSG00000275264 | downstream | 4757 |
| chr8 | 133723009 | 133723262 | ENSG00000253970 | upstream | 31549 |

| | | | | | |
|---|---|---|---|---|---|
| chr9 | 4866399 | 4866619 | ENSG00000228165 | upstream | 16026 |
| chr9 | 77596942 | 77597156 | ENSG00000228248 | downstream | 16314 |
| chr9 | 92152017 | 92152222 | ENSG00000236717 | inside | 1635 |
| chr9 | 94783014 | 94783390 | ENSG00000252153 | upstream | 26572 |
| chr9 | 123327904 | 123328093 | ENSG00000148204 | upstream | 28077 |
| chr9 | 126643032 | 126643221 | ENSG00000238010 | downstream | 1739 |
| chr9 | 128022298 | 128022530 | ENSG00000230848 | downstream | 16605 |
| chr9 | 128026919 | 128027189 | ENSG00000230848 | downstream | 11946 |
| chr9 | 128047346 | 128047521 | ENSG00000230848 | inside | 8211 |
| chr9 | 128298849 | 128299159 | ENSG00000272960 | downstream | 6002 |
| chr9 | 135392523 | 135392850 | ENSG00000235572 | downstream | 38303 |
| chr9 | 137513983 | 137514313 | ENSG00000263933 | upstream | 31435 |
| chr10 | 5496640 | 5496845 | ENSG00000178372 | downstream | 1852 |
| chr10 | 5660341 | 5660520 | ENSG00000196372 | inside | 6075 |
| chr10 | 17445789 | 17445995 | ENSG00000148488 | inside | 8335 |
| chr10 | 19232468 | 19232650 | ENSG00000227734 | downstream | 57573 |
| chr10 | 43026466 | 43026706 | ENSG00000263795 | downstream | 28785 |
| chr10 | 44173631 | 44173858 | ENSG00000237590 | downstream | 85744 |
| chr10 | 46048159 | 46048366 | ENSG00000263639 | upstream | 1890 |
| chr10 | 49883946 | 49884134 | ENSG00000230166 | upstream | 25479 |
| chr10 | 88385640 | 88385853 | ENSG00000275015 | upstream | 126268 |
| chr10 | 102710759 | 102710990 | ENSG00000138175 | inside | 3417 |
| chr10 | 125624871 | 125625069 | ENSG00000214297 | downstream | 41806 |
| chr10 | 133278859 | 133279036 | ENSG00000151651 | upstream | 1991 |
| chr11 | 1796612 | 1796866 | ENSG00000230980 | downstream | 3742 |
| chr11 | 2193367 | 2193565 | ENSG00000265258 | downstream | 20229 |
| chr11 | 69515342 | 69515582 | ENSG00000255774 | upstream | 35402 |
| chr11 | 69891604 | 69891898 | ENSG00000260348 | downstream | 17286 |

| | | | | | |
|---|---|---|---|---|---|
| chr11 | 70805902 | 70806189 | ENSG00000171671 | upstream | 56601 |
| chr11 | 71481506 | 71481729 | ENSG00000278349 | downstream | 7938 |
| chr11 | 72786500 | 72786727 | ENSG00000265064 | downstream | 2908 |
| chr11 | 76772651 | 76772868 | ENSG00000254632 | upstream | 4428 |
| chr11 | 78156424 | 78156650 | ENSG00000246174 | inside | 16653 |
| chr11 | 118057947 | 118058127 | ENSG00000272075 | upstream | 9110 |
| chr12 | 6364005 | 6364193 | ENSG00000111321 | upstream | 10852 |
| chr12 | 33587903 | 33588127 | ENSG00000212475 | downstream | 11703 |
| chr12 | 46493792 | 46493995 | ENSG00000272369 | upstream | 43507 |
| chr12 | 52148782 | 52149026 | ENSG00000265804 | downstream | 34770 |
| chr12 | 52203599 | 52203802 | ENSG00000257137 | downstream | 1782 |
| chr12 | 52972086 | 52972299 | ENSG00000265039 | downstream | 20291 |
| chr12 | 131860850 | 131861101 | ENSG00000256955 | inside | 3430 |
| chr13 | 38684839 | 38685138 | ENSG00000150893 | upstream | 1991 |
| chr13 | 98650387 | 98650585 | ENSG00000233662 | upstream | 9148 |
| chr13 | 112913573 | 112913780 | ENSG00000126217 | inside | 19195 |
| chr14 | 49992979 | 49993180 | ENSG00000279627 | inside | 675 |
| chr14 | 73756653 | 73756861 | ENSG00000264741 | upstream | 1886 |
| chr14 | 73784839 | 73785014 | ENSG00000259065 | upstream | 2346 |
| chr14 | 77400253 | 77400590 | ENSG00000280308 | upstream | 15808 |
| chr14 | 93017433 | 93017652 | ENSG00000258730 | upstream | 49800 |
| chr14 | 95056504 | 95056759 | ENSG00000258866 | downstream | 5551 |
| chr14 | 103693517 | 103693737 | ENSG00000269940 | upstream | 823 |
| chr15 | 39668539 | 39668793 | ENSG00000259279 | upstream | 74308 |
| chr15 | 71096700 | 71096947 | ENSG00000187720 | upstream | 5 |
| chr15 | 71104801 | 71105007 | ENSG00000187720 | inside | 7849 |
| chr15 | 73710228 | 73710414 | ENSG00000276807 | downstream | 19634 |
| chr15 | 74817462 | 74817715 | ENSG00000261606 | inside | 1239 |

| | | | | | |
|---|---|---|---|---|---|
| chr15 | 80679490 | 80679694 | ENSG00000136379 | overlapStart | 10 |
| chr15 | 89441379 | 89441566 | ENSG00000140519 | downstream | 29832 |
| chr15 | 95911741 | 95911936 | ENSG00000275443 | upstream | 78646 |
| chr15 | 96103339 | 96103541 | ENSG00000280048 | downstream | 9087 |
| chr16 | 327167 | 327421 | ENSG00000103126 | inside | 25252 |
| chr16 | 4152291 | 4152608 | ENSG00000262471 | downstream | 21471 |
| chr16 | 23137976 | 23138211 | ENSG00000103404 | inside | 11059 |
| chr16 | 23540350 | 23540636 | ENSG00000260247 | downstream | 2794 |
| chr16 | 68745774 | 68745953 | ENSG00000200558 | downstream | 3186 |
| chr16 | 69324652 | 69324843 | ENSG00000258429 | downstream | 3778 |
| chr16 | 70136707 | 70136937 | ENSG00000090857 | inside | 23081 |
| chr16 | 70390840 | 70391033 | ENSG00000260111 | inside | 8469 |
| chr16 | 72929501 | 72929707 | ENSG00000279757 | downstream | 43667 |
| chr16 | 75029907 | 75030118 | ENSG00000103091 | upstream | 29734 |
| chr16 | 81275815 | 81276079 | ENSG00000280182 | upstream | 11724 |
| chr16 | 83946915 | 83947134 | ENSG00000140961 | upstream | 1148 |
| chr16 | 88927159 | 88927345 | ENSG00000261226 | downstream | 9434 |
| chr17 | 40322341 | 40322612 | ENSG00000131759 | inside | 13149 |
| chr17 | 40576173 | 40576461 | ENSG00000126353 | upstream | 10701 |
| chr17 | 45043269 | 45043446 | ENSG00000136448 | upstream | 8164 |
| chr17 | 50944405 | 50944796 | ENSG00000247011 | overlapStart | 83 |
| chr17 | 58639731 | 58639989 | ENSG00000212195 | upstream | 7895 |
| chr17 | 61723167 | 61723572 | ENSG00000253506 | upstream | 131965 |
| chr17 | 68328998 | 68329179 | ENSG00000108932 | upstream | 37731 |
| chr17 | 72618037 | 72618240 | ENSG00000243514 | upstream | 8991 |
| chr17 | 74760697 | 74760983 | ENSG00000266036 | upstream | 11785 |
| chr17 | 74769820 | 74770055 | ENSG00000109065 | downstream | 492 |
| chr17 | 78421220 | 78421565 | ENSG00000200063 | downstream | 21157 |

| | | | | | |
|---|---|---|---|---|---|
| chr18 | 13539891 | 13540082 | ENSG00000272746 | downstream | 13203 |
| chr18 | 21330224 | 21330420 | ENSG00000265948 | downstream | 12429 |
| chr18 | 48990953 | 48991155 | ENSG00000280335 | upstream | 21951 |
| chr19 | 1181751 | 1182005 | ENSG00000118046 | inside | 4193 |
| chr19 | 1513239 | 1513421 | ENSG00000185761 | inside | 183 |
| chr19 | 18447033 | 18447265 | ENSG00000279262 | upstream | 1010 |
| chr19 | 33629591 | 33629772 | ENSG00000124302 | inside | 7636 |
| chr19 | 38299300 | 38299511 | ENSG00000167644 | upstream | 4650 |
| chr19 | 38714506 | 38714717 | ENSG00000267375 | upstream | 20900 |
| chr20 | 19283596 | 19283788 | ENSG00000179447 | inside | 808 |
| chr20 | 48075939 | 48076239 | ENSG00000276923 | downstream | 1751 |
| chr20 | 48158104 | 48158292 | ENSG00000276815 | downstream | 67886 |
| chr20 | 50727418 | 50727762 | ENSG00000124171 | upstream | 3782 |
| chr20 | 50764171 | 50764416 | ENSG00000124243 | upstream | 30478 |
| chr20 | 51387625 | 51387812 | ENSG00000263645 | upstream | 10304 |
| chr20 | 53661865 | 53662088 | ENSG00000238468 | downstream | 6609 |
| chr20 | 53725222 | 53725409 | ENSG00000238468 | upstream | 56464 |
| chr20 | 54195377 | 54195615 | ENSG00000101132 | upstream | 12232 |
| chr20 | 56735319 | 56735548 | ENSG00000231604 | downstream | 3859 |
| chr20 | 56758830 | 56759125 | ENSG00000207158 | downstream | 26856 |
| chr20 | 59988152 | 59988366 | ENSG00000124215 | inside | 29725 |
| chr21 | 31529496 | 31529673 | ENSG00000237594 | upstream | 29572 |
| chr21 | 42366609 | 42366957 | ENSG00000160182 | upstream | 15 |
| chr21 | 42376355 | 42376642 | ENSG00000160182 | upstream | 9761 |
| chr21 | 45330259 | 45330546 | ENSG00000229382 | upstream | 6161 |
| chr21 | 46292577 | 46293284 | ENSG00000182362 | inside | 4467 |
| chr21 | 46294298 | 46294590 | ENSG00000182362 | inside | 3161 |
| chr22 | 28813755 | 28814030 | ENSG00000272858 | upstream | 884 |

### Table S3α Gained ER binding sites

| Chromosome | Start | End | Feature | Location | Distance (bp) |
|---|---|---|---|---|---|
| chr1 | 107688502 | 107688700 | ENSG00000224550 | downstream | 183054 |
| chr1 | 107756462 | 107756674 | ENSG00000275455 | upstream | 140549 |
| chr1 | 113812315 | 113812588 | ENSG00000231128 | overlapStart | 64 |
| chr1 | 172746436 | 172746647 | ENSG00000229785 | upstream | 1913 |
| chr1 | 173868284 | 173868571 | ENSG00000185278 | inside | 202 |
| chr1 | 176752585 | 176752843 | ENSG00000237514 | upstream | 135799 |
| chr1 | 180167962 | 180168171 | ENSG00000116260 | inside | 13128 |
| chr1 | 181022548 | 181023230 | ENSG00000135823 | overlapStart | 109 |
| chr1 | 185645554 | 185645864 | ENSG00000261729 | upstream | 9138 |
| chr1 | 186152804 | 186153261 | ENSG00000224691 | downstream | 23553 |
| chr1 | 188432511 | 188432717 | ENSG00000225006 | upstream | 75821 |
| chr1 | 191153132 | 191153513 | ENSG00000261642 | inside | 1121 |
| chr1 | 200150699 | 200150956 | ENSG00000229220 | upstream | 2420 |
| chr1 | 205122017 | 205122304 | ENSG00000117222 | upstream | 2 |
| chr2 | 10233 | 10452 | ENSG00000184731 | downstream | 28362 |
| chr2 | 20071741 | 20072135 | ENSG00000223734 | downstream | 17245 |
| chr2 | 38076093 | 38076926 | ENSG00000232973 | inside | 445 |
| chr2 | 38690697 | 38691078 | ENSG00000232518 | upstream | 19276 |
| chr2 | 43463263 | 43463515 | ENSG00000264071 | upstream | 28517 |
| chr2 | 70616316 | 70616556 | ENSG00000163235 | upstream | 62123 |
| chr2 | 88181019 | 88181590 | ENSG00000144115 | inside | 5046 |
| chr2 | 134083462 | 134083789 | ENSG00000152127 | upstream | 36194 |
| chr2 | 139245261 | 139245613 | ENSG00000226939 | upstream | 120552 |
| chr2 | 214378717 | 214379091 | ENSG00000174453 | upstream | 31974 |

| | | | | | |
|---|---|---|---|---|---|
| chr2 | 215216955 | 215217179 | ENSG00000237571 | upstream | 57813 |
| chr2 | 215982550 | 215982933 | ENSG00000226276 | upstream | 40831 |
| chr2 | 217092662 | 217092912 | ENSG00000251849 | upstream | 83586 |
| chr2 | 226151187 | 226151425 | ENSG00000235070 | downstream | 28619 |
| chr2 | 236824952 | 236825181 | ENSG00000232328 | downstream | 70589 |
| chr2 | 237502454 | 237502717 | ENSG00000277842 | upstream | 8214 |
| chr3 | 4978425 | 4979079 | ENSG00000134107 | upstream | 37 |
| chr3 | 125916135 | 125916404 | ENSG00000171084 | upstream | 216 |
| chr3 | 138609262 | 138609711 | ENSG00000158234 | inside | 656 |
| chr3 | 158723252 | 158723520 | ENSG00000118855 | upstream | 8678 |
| chr3 | 174955657 | 174955861 | ENSG00000230497 | downstream | 103500 |
| chr3 | 175127135 | 175127500 | ENSG00000230292 | upstream | 11893 |
| chr3 | 175136676 | 175136913 | ENSG00000230292 | upstream | 21434 |
| chr4 | 3954973 | 3955232 | ENSG00000251669 | inside | 187 |
| chr4 | 3954973 | 3955232 | ENSG00000253917 | inside | 187 |
| chr4 | 3955472 | 3955892 | ENSG00000251669 | upstream | 53 |
| chr4 | 3955472 | 3955892 | ENSG00000253917 | upstream | 53 |
| chr4 | 88318986 | 88319228 | ENSG00000207480 | downstream | 10948 |
| chr4 | 88335030 | 88335258 | ENSG00000207480 | upstream | 4752 |
| chr4 | 105126504 | 105126701 | ENSG00000251259 | downstream | 10579 |
| chr4 | 114326430 | 114326658 | ENSG00000248716 | downstream | 222205 |
| chr4 | 140810785 | 140811056 | ENSG00000222613 | upstream | 31644 |
| chr4 | 142615364 | 142615633 | ENSG00000249806 | inside | 45317 |
| chr5 | 49497 | 49789 | ENSG00000250020 | upstream | 8409 |
| chr5 | 17306048 | 17306482 | ENSG00000185296 | overlapStart | 101 |
| chr5 | 71748521 | 71748789 | ENSG00000278824 | downstream | 5589 |
| chr5 | 76699566 | 76699938 | ENSG00000277188 | upstream | 12257 |
| chr5 | 129355259 | 129355480 | ENSG00000265691 | downstream | 41582 |

| | | | | | |
|---|---|---|---|---|---|
| chr5 | 147930482 | 147930686 | ENSG00000250346 | downstream | 7061 |
| chr6 | 21738659 | 21739119 | ENSG00000272168 | inside | 73887 |
| chr6 | 37697767 | 37698332 | ENSG00000112139 | inside | 974 |
| chr6 | 67086533 | 67086824 | ENSG00000266073 | downstream | 62685 |
| chr6 | 73520999 | 73521230 | ENSG00000229862 | upstream | 2388 |
| chr6 | 82570085 | 82570355 | ENSG00000146242 | downstream | 199257 |
| chr6 | 87574255 | 87574504 | ENSG00000146282 | inside | 15499 |
| chr6 | 122620965 | 122621352 | ENSG00000272472 | upstream | 22036 |
| chr6 | 122696980 | 122697209 | ENSG00000240606 | upstream | 48071 |
| chr6 | 151658105 | 151658419 | ENSG00000091831 | inside | 1414 |
| chr6 | 151668982 | 151669253 | ENSG00000091831 | inside | 12291 |
| chr7 | 23273231 | 23273439 | ENSG00000156928 | upstream | 25300 |
| chr7 | 23334717 | 23334949 | ENSG00000232627 | upstream | 30669 |
| chr7 | 34037057 | 34037290 | ENSG00000164619 | inside | 118582 |
| chr7 | 34069707 | 34069904 | ENSG00000236212 | upstream | 139622 |
| chr7 | 101973172 | 101973370 | ENSG00000272219 | downstream | 11278 |
| chr7 | 120717279 | 120717513 | ENSG00000231295 | downstream | 29225 |
| chr7 | 159013236 | 159013688 | ENSG00000231419 | inside | 6714 |
| chr8 | 33245099 | 33245358 | ENSG00000253993 | upstream | 17952 |
| chr8 | 37144873 | 37145113 | ENSG00000275998 | upstream | 28716 |
| chr8 | 66522552 | 66522752 | ENSG00000206949 | upstream | 21241 |
| chr8 | 66713977 | 66714328 | ENSG00000104205 | inside | 1559 |
| chr8 | 78530378 | 78530727 | ENSG00000171033 | inside | 14239 |
| chr8 | 90754309 | 90754561 | ENSG00000123119 | upstream | 36989 |
| chr8 | 96945865 | 96946114 | ENSG00000272249 | downstream | 186721 |
| chr8 | 100310060 | 100310486 | ENSG00000253217 | upstream | 27109 |
| chr8 | 102653875 | 102654237 | ENSG00000155090 | inside | 1665 |
| chr8 | 115838444 | 115838697 | ENSG00000104447 | upstream | 28771 |

| | | | | | |
|---|---|---|---|---|---|
| chr8 | 115998303 | 115998591 | ENSG00000199450 | downstream | 24095 |
| chr8 | 118878632 | 118878904 | ENSG00000164761 | downstream | 44653 |
| chr8 | 121145570 | 121145820 | ENSG00000271219 | upstream | 31651 |
| chr8 | 127859778 | 127860054 | ENSG00000278324 | upstream | 30572 |
| chr8 | 133216514 | 133216731 | ENSG00000270132 | upstream | 12325 |
| chr9 | 70419775 | 70420150 | ENSG00000119138 | upstream | 5151 |
| chr9 | 80913206 | 80913420 | ENSG00000226798 | upstream | 41911 |
| chr9 | 106863871 | 106864117 | ENSG00000148143 | inside | 774 |
| chr9 | 127955784 | 127956003 | ENSG00000136908 | upstream | 17300 |
| chr9 | 129488720 | 129488919 | ENSG00000204054 | inside | 5269 |
| chr9 | 133563733 | 133564751 | ENSG00000196990 | downstream | 13664 |
| chr10 | 1049157 | 1049382 | ENSG00000067064 | overlapStart | 13 |
| chr10 | 9188219 | 9188778 | ENSG00000230014 | upstream | 87055 |
| chr10 | 103168446 | 103168687 | ENSG00000276694 | upstream | 3250 |
| chr10 | 131966899 | 131967109 | ENSG00000277959 | upstream | 4093 |
| chr10 | 131969166 | 131969616 | ENSG00000277959 | upstream | 1586 |
| chr11 | 3422697 | 3422939 | ENSG00000166492 | upstream | 201 |
| chr11 | 67805577 | 67805775 | ENSG00000160172 | upstream | 241 |
| chr11 | 69122852 | 69123095 | ENSG00000260895 | downstream | 13758 |
| chr11 | 69654515 | 69654934 | ENSG00000110092 | downstream | 41 |
| chr11 | 71787087 | 71787322 | ENSG00000158483 | upstream | 188 |
| chr11 | 76788521 | 76788866 | ENSG00000255100 | upstream | 5459 |
| chr11 | 85828179 | 85828562 | ENSG00000137501 | upstream | 17020 |
| chr12 | 12872431 | 12872759 | ENSG00000234498 | upstream | 2740 |
| chr12 | 33609629 | 33609834 | ENSG00000212475 | downstream | 33429 |
| chr12 | 39443233 | 39443482 | ENSG00000139116 | overlapStart | 92 |
| chr12 | 45596374 | 45596691 | ENSG00000257657 | inside | 13929 |
| chr12 | 53157996 | 53158215 | ENSG00000257808 | upstream | 1371 |

| | | | | | |
|---|---|---|---|---|---|
| chr12 | 53825423 | 53825652 | ENSG00000253053 | downstream | 8944 |
| chr12 | 54892183 | 54892403 | ENSG00000172551 | downstream | 33790 |
| chr12 | 57088666 | 57088909 | ENSG00000166886 | overlapStart | 15 |
| chr12 | 75236287 | 75236501 | ENSG00000254451 | inside | 1547 |
| chr14 | 35403822 | 35404255 | ENSG00000100906 | inside | 494 |
| chr14 | 40677050 | 40677339 | ENSG00000251363 | upstream | 277559 |
| chr14 | 59361251 | 59361646 | ENSG00000252869 | upstream | 38377 |
| chr15 | 33068011 | 33068216 | ENSG00000279958 | downstream | 37542 |
| chr15 | 67840045 | 67840293 | ENSG00000206625 | overlapStart | 0 |
| chr15 | 86864105 | 86864324 | ENSG00000273540 | upstream | 50119 |
| chr15 | 88638685 | 88638905 | ENSG00000172183 | inside | 2532 |
| chr15 | 89115627 | 89115845 | ENSG00000239151 | upstream | 4934 |
| chr15 | 92904276 | 92904518 | ENSG00000264173 | includeFeature | 43 |
| chr15 | 95247161 | 95247511 | ENSG00000273923 | upstream | 21297 |
| chr15 | 96336968 | 96337231 | ENSG00000222651 | downstream | 3661 |
| chr16 | 19598 | 19901 | ENSG00000234769 | upstream | 146 |
| chr16 | 9047736 | 9047988 | ENSG00000261392 | upstream | 20566 |
| chr16 | 30065286 | 30065595 | ENSG00000274904 | upstream | 461 |
| chr16 | 68234354 | 68234615 | ENSG00000276151 | upstream | 855 |
| chr16 | 77604073 | 77604293 | ENSG00000261154 | inside | 6388 |
| chr16 | 88650799 | 88651819 | ENSG00000051523 | overlapStart | 353 |
| chr16 | 88652628 | 88653682 | ENSG00000051523 | upstream | 1476 |
| chr16 | 88653897 | 88654213 | ENSG00000051523 | upstream | 2745 |
| chr17 | 41536024 | 41536337 | ENSG00000171345 | upstream | 7716 |
| chr17 | 59837417 | 59837946 | ENSG00000199004 | upstream | 3320 |
| chr17 | 64506366 | 64506652 | ENSG00000258890 | overlapStart | 64 |
| chr17 | 65243744 | 65243984 | ENSG00000265883 | upstream | 58262 |
| chr17 | 75549401 | 75549690 | ENSG00000073350 | inside | 24321 |

| | | | | | |
|---|---|---|---|---|---|
| chr17 | 82092902 | 82097065 | ENSG00000169710 | inside | 1267 |
| chr17 | 82614043 | 82614484 | ENSG00000261845 | upstream | 9865 |
| chr18 | 43876827 | 43877030 | ENSG00000202250 | upstream | 44562 |
| chr18 | 45802729 | 45802996 | ENSG00000267558 | downstream | 14062 |
| chr18 | 60053882 | 60054141 | ENSG00000252555 | downstream | 35151 |
| chr19 | 1170358 | 1170567 | ENSG00000064932 | inside | 3716 |
| chr19 | 18281761 | 18282047 | ENSG00000267959 | upstream | 30 |
| chr19 | 42242758 | 42243176 | ENSG00000279539 | overlapStart | 17 |
| chr20 | 53805933 | 53806163 | ENSG00000225563 | upstream | 4615 |
| chr20 | 54149160 | 54149361 | ENSG00000019186 | downstream | 4088 |
| chr20 | 63101534 | 63101765 | ENSG00000274915 | upstream | 377 |
| chr20 | 64286740 | 64287339 | ENSG00000149656 | upstream | 3046 |
| chr21 | 21238955 | 21239193 | ENSG00000226771 | upstream | 12126 |
| chr21 | 37267892 | 37268174 | ENSG00000157538 | overlapStart | 27 |
| chr21 | 40340426 | 40340645 | ENSG00000235123 | upstream | 42438 |
| chr21 | 46134261 | 46134708 | ENSG00000237338 | upstream | 16906 |
| chr22 | 37328907 | 37329159 | ENSG00000237862 | downstream | 23031 |
| chrX | 16662400 | 16662673 | ENSG00000169906 | downstream | 7730 |
| chrX | 116285496 | 116285825 | ENSG00000180772 | downstream | 110524 |
| chrX | 131493185 | 131493396 | ENSG00000228450 | upstream | 299 |
| chrX | 152915532 | 152915742 | ENSG00000147394 | inside | 1090 |

Table S4 RNA-seq MCF7 Fulvestrant - Vehicle (adjusted p-value < 0.05)

Table S5 RNA-seq T47D Fulvestrant - Vehicle (adjusted p-value < 0.05)

Table S6 RNA-seq MCF7 Hypoxia - Normoxia (adjusted p-value < 0.05)

Table S7 RNA-seq T47D Hypoxia - Normoxia (adjusted p-value < 0.05)

Table S8 RNA-seq MCF7 Hypoxia - Normoxia with Fulvestrant (adjusted P-value < 0.05)

Table S9 RNA-seq T47D Hypoxia - Normoxia with Fulvestrant (adjusted P-value < 0.05)
